# Open and Hybridized: The Topology of Chloroplast Barrel Proteins

**DOI:** 10.64898/2026.07.15.738633

**Authors:** Emily Proctor, Daniel Montezano, Matthew M. Copeland, Joanna S. G. Slusky

## Abstract

Despite their central role in metabolite exchange, lipid trafficking, and protein import, chloroplast outer envelope beta-barrel proteins lack a dedicated comprehensive sequence database spanning many plant proteomes. Here we present the database COB (chloroplast outer-envelope beta-barrel), consisting of 16,586 beta-barrel sequences which we organize across ten protein categories and an uncharacterized group. COB was constructed using a machine learning classifier that identifies chloroplast beta-barrels based on features derived from evolutionary protein contact maps. Analysis of COB reveals that Streptophyta have more barrels overall and use a greater variety of solute transporters than Chlorophyta. Furthermore, we find considerable structural diversity across OEP categories, including variation in beta-strand count and a high prevalence of open barrel conformations not observed in bacterial outer membrane proteins. Structure predictions for *Arabidopsis thaliana* outer envelope proteins identified candidate hybrid barrel assemblies, with TOC159 family members emerging as universal interaction partners. We also report single-chain multi-barrel domain architectures in the chloroplast outer envelope, a topology previously described only in Gram-negative bacteria. Finally, we find chloroplast membrane barrels have more open topologies and shorter strands than bacterial membrane barrels. COB provides a comprehensive sequence resource for chloroplast outer envelope beta-barrels and establishes a foundation for investigating the evolution, structure, and function of the chloroplast outer envelope.

## Introduction

The chloroplast outer envelope carries the structural signatures of its evolutionary origin: it is what remains of a free-living cyanobacterium’s outer membrane, with features retained across more than a billion years of endosymbiosis even as the organelle’s genome and most of its proteins were absorbed into the eukaryotic host [1–3]. Since its absorption the chloroplast outer envelope has been pressed into service well beyond what any bacterial membrane was ever built to do. It is the sole conduit for the metabolites that link chloroplast metabolism to the rest of the cell, for the lipids that build and remodel its own membranes, for the ions and small solutes that photosynthesis depends on turning over continuously, and for the proteins — nearly all of them, since most of the ancestral genome moved to the host nucleus — that must be delivered into an organelle they, evolutionarily, never left [4, 5] [6–8]. All of this traffic is carried out not by transporters built from scratch, but by a family of proteins whose own evolutionary history is, if anything, even more distinctive: the beta-barrels.

The barrels embedded in the outer envelope are a structural class found only in the outer membranes (OM) of bacteria, mitochondria, and chloroplasts [9]. The chloroplast outer envelope contains a set of defined outer envelope protein (OEP) categories that can be broadly grouped by function: solute transporters (OEP21, OEP24, OEP37, OEP40), lipid transporters (TGD4 and LPTD1), Omp85 superfamily members (OEP80, SP2/P39, TOC75), and the TOC159 receptor family (TOC159, TOC132, TOC120, and TOC90). The four functional groups differ considerably in structure and function.

### Solute transport

The solute transporter group mediates passage of a diverse set of molecules across the outer envelope. OEP21 is an ATP-regulated, anion-selective channel. It serves as the exit pore for triose phosphates (TPs), 3-phosphoglycerate, and inorganic phosphate (Pi), which are central intermediates of photosynthetic carbon fixation [10–12]. OEP24 serves a complementary role, though with broader substrate specificity, including TPs, dicarboxylic acids, amino acids, sugars, ATP, and Pi [13, 14]. Although both channels participate in TP export, their relative abundance differs between plant species. OEP21 is prominent in C3 plants, while OEP24 is more abundant in C4 plants where its broader conductance and lower selectivity are hypothesized to better support higher metabolite rates [15]. In contrast, OEP37 is a cation-selective channel whose substrate and primary role remain poorly defined [16], while OEP40 selectively facilitates glucose transport and its phosphorylated derivatives [17]. The diversity shown across these channels reflects the complexity and specificity of metabolite exchange at the chloroplast OE.

### Lipid transport

The lipid transporter group is represented by TGD4, which is a phosphatidic acid binding beta-barrel [18, 19]. It is distantly related to the bacterial outer membrane protein LptD, which transports lipopolysaccharide across the outer membrane of Gram-negative bacteria [20]. Leaf glycerolipid composition in *Arabidopsis* is strongly enriched in chloroplast associated galactolipids, with mono-(MGDG) and digalactosyldiacylglycerol (DGDG) together accounting for about 60% of leaf glycerolipids [21]. Chloroplasts synthesize glycerolipids through two routes, a prokaryotic pathway retained entirely within the chloroplast, and a eukaryotic pathway in which fatty acids are exported to the ER for lipid assembly and subsequently reimported into the chloroplast. TGD4 functions in the eukaryotic pathway, where it binds phosphatidic acid at the outer envelope and facilitates reimport of ER-derived lipid precursors into the chloroplast. In *Arabidopsis*, a second LptD-family protein, LPTD1, is predicted to function as a lipid transporter with a particular role in glycerolipid remodeling under phosphate starvation [22].

### Protein movement

The Omp85 superfamily is a conserved family of outer membrane proteins. Bacterial BamA in this superfamily inserts beta-barrels into the bacterial outer membrane [23]. The chloroplast outer envelope contains multiple members of this superfamily, each serving distinct roles related to outer membrane biogenesis. OEP80 (TOC75-V) is thought to perform a function analogous to BamA [24]. Another Omp85 protein, SP2/P39, is evolutionarily related to the OEP80 family, but lacks the characteristic N-terminal POTRA domains. It functions as the core channel of the chloroplast-associated protein degradation complex (CHLORAD), mediating extraction of ubiquitinated OEPs for degradation [25]. Lastly, TOC75 is also a member of the Omp85 superfamily, and serves as the core channel of the TOC complex through which preproteins are translocated into the chloroplast [26].

### Receptors

The TOC159 family are the GTPase receptor components of the TOC complex, responsible for recognizing preproteins at the chloroplast surface. This is encoded by a small gene family with four homologous members, TOC159, TOC132, TOC120, and TOC90. Each shares a tripartite domain architecture consisting of an N-terminal acidic domain, a central GTPase domain, and a C-terminal membrane anchor domain that forms a beta-barrel embedded in the outer envelope. Although, the acidic domain is greatly truncated in TOC90 and sequence identity varies across paralogs [27–29]. Beyond their receptor function, the C-terminal beta-barrel domain also contributes structurally to the translocation pore itself. The TOC75 and TOC159 family members come together to form a continuous hybrid beta-barrel pore through which preproteins are translocated in the TOC complex [30, 31]. In this structure, strands from each protein contribute to a single, large continuous barrel. These paralogs assemble into compositionally distinct TOC complexes with TOC75. TOC159, and to a lesser extent TOC90, preferentially facilitate import of photosynthetic preproteins, while TOC132 and TOC120 mediate import of housekeeping proteins [27–29, 32]. Together, TOC159 family members and TOC75 form the core machinery through which the vast majority of chloroplast preproteins are recognized and imported — and their hybrid pore raises a broader question: whether barrel-sharing of this kind is a peculiarity of the TOC complex, or a more general feature of chloroplast outer envelope architecture that has simply gone undetected elsewhere.

Efforts to catalog chloroplast outer envelope beta-barrels at scale have progressed through both proteomic and computational approaches, each advancing the field only partway. Mass spectrometry surveys of spinach and Arabidopsis envelope fractions [33–35]., refined over successive studies through improved sensitivity, cross-species comparison, and curated localization databases [36–38], identified hundreds of envelope-associated proteins, but consolidated reviews still place the number of characterized chloroplast beta-barrel genes in Arabidopsis at only about 18 (Supp. Table 1) [39, 40] — a reflection of how few of those candidates have been confirmed as beta-barrels rather than merely surveyed as envelope-associated. Computational prediction has closed part of this gap but introduced its own limitations. An early attempt adapted a bacterial beta-barrel scoring algorithm to the Arabidopsis genome but struggled to distinguish barrels from soluble proteins and was restricted to model organisms [41, 42].

**Table 1.**
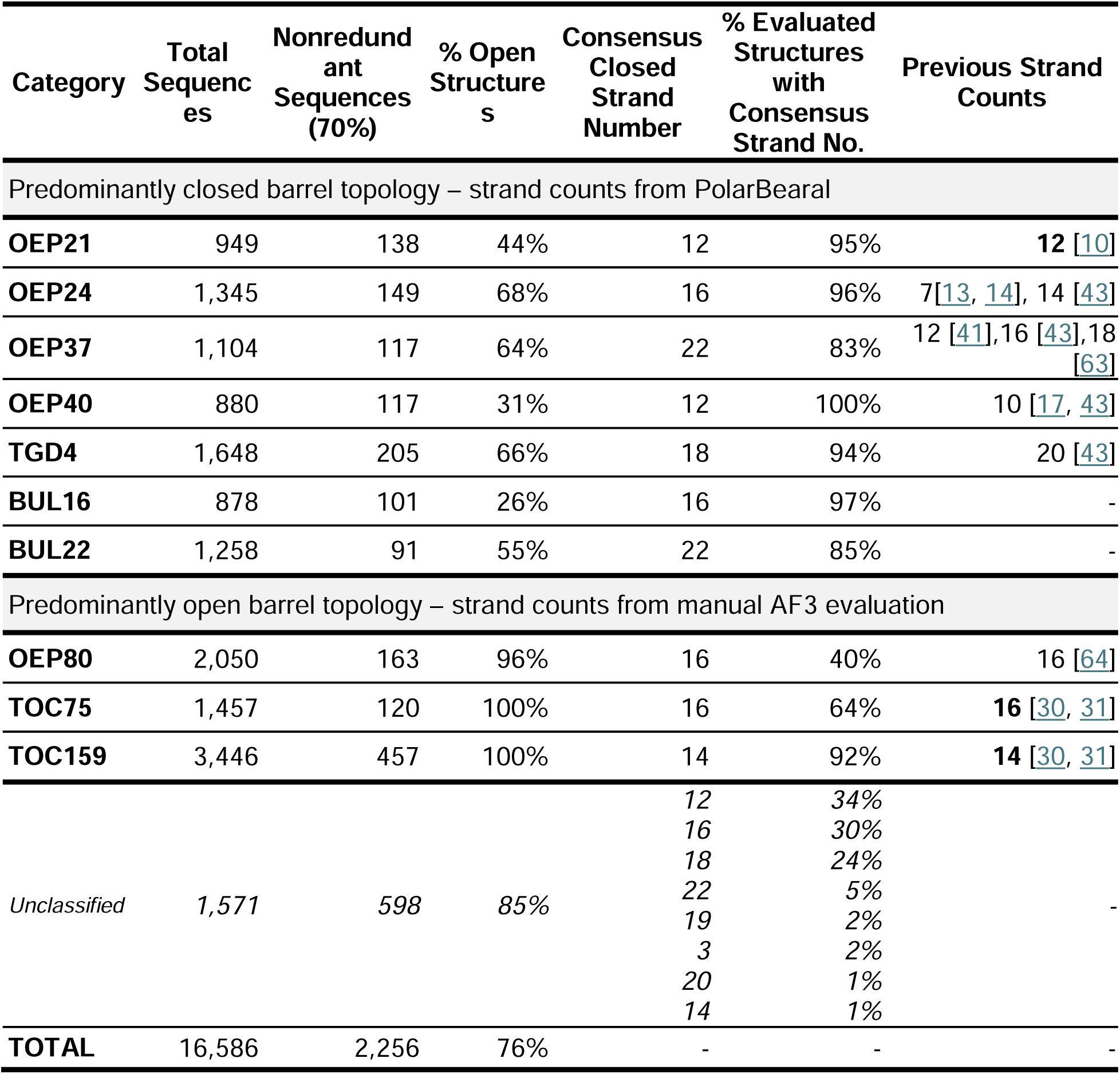
Structural Classification for OEP Beta-Barrel Categories in COB. Strand counts were assigned using PolarBearal for predominantly closed barrel categories, and by manual evaluation of AF3 predicted structures for predominantly open barrel categories. For closed barrel categories, % Open Structures reflects the fraction of non-redundant sequences for which PolarBearal failed to return a prediction. For open barrel categories, % Open Structures was determined by manual evaluation of AF3 predicted structures. Consensus Closed Strand Number indicates the most frequently observed strand count, and % of Evaluated Structures indicates the percentage matching that consensus. Previous strand counts are values reported in prior literature, with bold indicating counts determined from experimental structures.

More recently, Roumia and co-authors built profile hidden Markov models (pHMMs) for eukaryotic beta-barrel families of chloroplasts and mitochondria [43] and detected approximately 2,000 chloroplast beta-barrels across eukaryotic reference proteomes [44] — an order-of-magnitude advance, but one still bounded by sequence similarity to previously characterized families. Divergent homologs and families not yet recognized as beta-barrels fall outside this net entirely: the TOC159 receptor family, for instance, was not yet known to form a beta-barrel and was absent from these searches, while conversely OEP23 [45] was included as a beta-barrel despite structural predictions indicating it does not adopt one, consistent with earlier observations [46]. Despite this progression, no comprehensive database dedicated to chloroplast beta-barrel proteins across diverse plant proteomes currently exists.

Here, we present a machine learning classifier that identifies chloroplast beta-barrels from features derived from evolutionary contact maps rather than full structure prediction, since the majority of chloroplast outer envelope proteins across plant proteomes still lack AlphaFold-predicted structures and large-scale 3D prediction remains computationally prohibitive at proteome scale [47]. Because beta-barrels’ regular hydrogen-bonding pattern between antiparallel strands leaves a distinctive signature in these 2D contact maps, this approach enables classification across entire plant proteomes at a scale full structure prediction cannot match; we then use AF structure prediction selectively, to validate high-confidence classifier calls rather than as a primary discovery tool.

Applying this classifier across Viridiplantae, we constructed COB, a database of 16,586 chloroplast outer envelope beta-barrel proteins spanning ten OEP categories — an over eight-fold expansion of the known chloroplast beta-barrel repertoire. Analysis of COB reveals that chloroplast barrels are, on average, more open and shorter-stranded than their bacterial counterparts; identifies candidate hybrid barrel assemblies beyond the TOC complex, with TOC159 family members emerging as recurring interaction partners; and uncovers single-chain multi-barrel domain architectures in the chloroplast outer envelope, a topology previously documented only in Gram-negative bacteria. Together, these findings establish COB as both a comprehensive sequence resource and a foundation for investigating how this bacterially derived protein family has been repurposed and diversified over a billion years of chloroplast evolution.

## Results

### COB classifier

Our chloroplast outer envelope beta-barrel classifier is adapted from a previously developed algorithm that successfully predicted beta-barrel proteins in the bacterial outer membrane [48]. We hypothesized that our bacterial barrel classifier could be extended to the evolutionarily related chloroplast outer envelope. Briefly, the three steps of our classifier are: generate an evolutionary contact map, contact map feature extraction, classification.

Evolutionary contact maps were generated by RaptorX [49]. Beta-barrel proteins generate a distinctive pattern of diagonal contacts in their contact maps that are perpendicular to the self-interaction diagonal. These slashes reflect regular hydrogen bonding between adjacent beta-strands (Fig. 1A). Five features are extracted from the contact maps, including the number of beta-hairpins, average hairpin length, hairpin spacing standard deviation, a closing contact confidence score, and the protein sequence length (Fig. 1B-C). These features serve as input to a random forest classifier (Fig. 1D). The classifier was trained on a dataset we curated of predicted chloroplast beta-barrels and non-barrel proteins (1,537DS, See Methods). The model achieved a balanced accuracy of 85.1%, precision of 95.1%, and a Matthews Correlation Coefficient (MCC) of 75.8% when tested against a 10% hold-out set, demonstrating reliable discrimination between barrel and non-barrel proteins while minimizing false positives.

**Figure 1.**
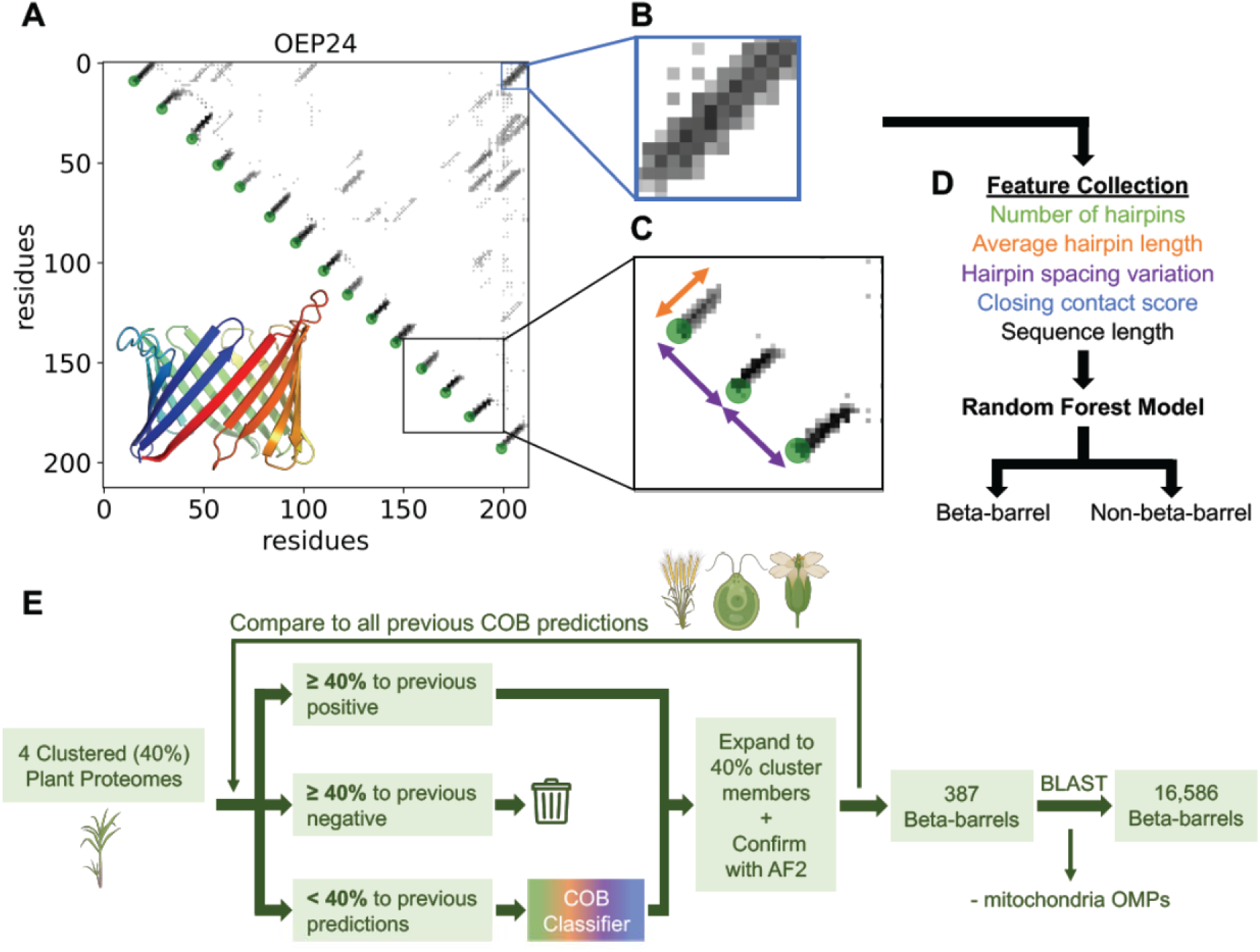
COB Classifier and COB Creation. (A) Contact map of OEP24 (NCBI: NP_175131.3), showing the characteristic pattern produced by interstrand hydrogen bonding. Maps are symmetrical over the diagonal, so we used the lower (triangular) half for the predicted structure of the same sequence. (B) Zoomed in view of the contact map highlighting the closing contact interaction between the first and last strand (blue). (C) Zoomed-in view of contact map illustrating individual beta-hairpin features, with arrows indicating average hairpin length (orange) and spacing (purple). (D) COB classification pipeline. Five features, including the number of beta-hairpins, average beta-hairpin length, beta-hairpin spacing standard deviation, closing contact score, and sequence length, are extracted from the contact map and provided as input to a random forest classifier, which outputs a binary prediction of beta-barrel or non-beta-barrel. (E) Workflow for COB construction. Proteomes from four Viridiplantae species (T. aestivum, C. reinhardtii, A. thaliana, and Saccharum hybrid cultivar R570) were clustered at 40% sequence identity. Representatives were compared to sequences previously predicted by the COB classifier. Those exceeding the 40% threshold took on the same COB prediction, while contact maps were generated for the remaining sequences and predicted with COB classifier. Predicted beta-barrels were expanded to include cluster members and validated with AlphaFold2 (AF2), ultimately yielding 387 chloroplast beta-barrel proteins. BLAST expansion against the NCBI nr database, followed by sequence similarity clustering, structural reclassification of uncharacterized sequences, and removing any mitochondria outer membrane proteins (OMPs), produced a final database of 16,586 chloroplast beta-barrels. (Plant images created with BioRender).

### COB construction

#### Barrel identification

To build a comprehensive database of chloroplast outer envelope beta-barrels, we first applied the COB classifier to four representative Viridiplantae proteomes to generate a high-confidence initial set. We then expanded that set through database searching, and finally classified the sequences into ten OEP categories.

To identify chloroplast beta-barrels across Viridiplantae, we applied our COB classifier iteratively against four representative proteomes (Fig. 1E). These proteomes were selected to represent diversity within Viridiplantae. We used *Arabidopsis thaliana* (thale cress) [50] as a well-characterized model organism, *Triticum aestivum* (bread wheat) [51] and *Chlamydomonas reinhardtii* (green algae) [52] as large NCBI proteomes for Streptophyta and Chlorophyta respectively, and *Saccharum officinarum x Saccharum spontaneum* hybrid cultivar R570 (sugarcane) [53] as an extra-large proteome. Contact map generation of whole proteomes is a process that still requires large computational time, so we first determined that sequences sharing ≥40% sequence similarity receive the same COB prediction. We found ≥40% sequence similarity yields >97% same classification (barrel or not barrel, Supp. Fig. 1). This allows us to bypass map generation for any sequence similar to a sequence previously predicted. Each proteome was therefore clustered at 40% sequence identity using MMseqs2 [54], and the COB classifier was applied to cluster representatives rather than all sequences.

We began with *T. aestivum,* for which all contact maps for the representative sequences were generated as no prior COB predictions existed. The same workflow was then applied iteratively to *C. reinhardtii, A. thaliana*, and *Saccharum* hybrid. For each successive proteome, representatives sharing ≥40% similarity with any sequence already predicted in a prior proteome inherited that prediction directly. Specifically, any representative sequence ≥40% similar to at least one barrel was assigned to be a barrel, whereas representative sequences only similar to non-barrel sequences were discarded. Barrel predictions were expanded to all members of their respective clusters.

As we searched whole proteomes, and outer membrane beta-barrels are found in both the chloroplast and mitochondrial outer membranes in eukaryotes, the COB classifier identified mitochondria annotated beta-barrels which were subsequently removed.

We discovered that some categories of chloroplast proteins (the families OEP40 and TOC159/132/120/90) were not identified by our COB algorithm. However, since they are known chloroplast barrels, we sought to include those in our database as well. Sequences from these families were therefore recovered using pHMM searches against all four proteomes. All candidate sequences from our workflow and the pHMM searches were manually validated using AlphaFold2 (AF2) structures to ensure membrane beta-barrel topology [47]. Together, the COB algorithm with the manual addition of OEP40 and TOC159/132/120/90 resulted in 344 unique sequences corresponding to 387 protein accessions, including 35 (*A. thaliana*), 9 (*C. reinhardtii*), 100 (*T. aestivum*), and 243 (*Saccharum* hybrid) beta-barrels, yielding an initial high-confidence set.

To expand this initial set, we BLAST searched the 344 unique sequences against the NCBI non-redundant (nr) database restricted to Viridiplantae, retrieving 30,286 unique candidate protein accessions [55, 56]. Of these, 20,244 sequences were sufficiently sequence similar (40%) to an existing barrel prediction to label them as barrels.

#### Categorizing barrels

To organize the 20,244 candidate barrel sequences into known chloroplast beta-barrel families, we classified them across OEP categories. The OEP categories are: OEP21, OEP24, OEP37, OEP40, OEP80, TOC75, TOC159, and TGD4. These categories group proteins by shared structural fold, and in some cases include homologs with distinct functions. OEP21 includes OEP21A and OEP21B, OEP24 includes OEP24A and OEP24B, TOC75 includes TOC75-III and TOC75-IV, TGD4 includes LPTD1, TOC159 includes TOC132, TOC120, and TOC90, and OEP80 (TOC75-V) includes SP2 (P39).

To categorize the proteins, we began with clustering the 20,244 sequences at 40% sequence identity using MMseqs2 [54]. Each cluster—composed of a representative sequence and its members—was assigned to the OEP category most represented across the NCBI annotations of all its members. This allowed us to classify large numbers of unannotated sequences by leveraging the consensus annotation of their cluster, rather than relying on individual sequence annotations alone. Clusters whose members could not reach a confident consensus assignment were placed in the Unclassified category. Clusters with predominantly mitochondrial beta-barrel annotations, including VDAC [57], Tom40 [58], or Sam50 [59], were removed from the dataset. After the initial sequence similarity clustering, some proteins in the Unclassified category had the fold of existing categories. Structural fold similarities can link proteins even after sequence similarity has been erased by evolution [60]. To better assign sequences remaining in the Unclassified category, we searched for fold similarities to representatives of each known OEP category using Foldseek [61]. Rather than classifying individual sequences, we applied a cluster-based approach using the MMseqs2 clusters and AlphaFold3 (AF3) structures (See Methods). Clusters whose members showed consensus structural similarity to mitochondrial barrels were removed, while those that could not be confidently assigned through either annotation or structure-based classification were retained in the Unclassified category. Finally, sequences less than 125 residues in length were removed from the dataset.

Together, these annotation and structure-based classification steps produced COB, a database of 16,586 chloroplast beta-barrel proteins. Along with our previously known OEP categories, we identified two prominent clusters within the Unclassified group that were structurally distinct from any known OEP type. Although these clusters passed our mitochondrial filtering steps, we cannot confidently assign them to the chloroplast outer envelope. They were therefore designated Barrel of Unknown Localization 16 (BUL16) and BUL22, and named for their consensus closed strand counts of 16 and 22, respectively. In total, COB spans ten categories, including eight established OEP families, BUL16, BUL22, and an Unclassified category.

Previous work in OMPdb used pHMMs to identify 1,933 chloroplast beta-barrels across eukaryotic reference proteomes [44]. COB represents an over eight-fold increase relative to this dataset. To understand where this increase comes from, we examined how COB sequences relate to what was available to OMPdb at the time of their search (Supp. Fig. 2). Sequences from Viridiplantae RefSeq proteomes available in 2020 or earlier, approximating the proteome space accessible to OMPdb, account for 7.7% of the increase in COB. An additional 11.5% of COB comes from RefSeq proteomes released after 2020, reflecting growth in publicly available reference plant proteomes since that time. OMPdb restricted its search to reference proteomes, while we searched all available Viridiplantae sequences in the NCBI non-redundant database. The use of the non-redundant database accounts for 47.1% of the increase. We were also able to incorporate chloroplast beta-barrel types not present in OMPdb. The TOC159 receptor family was not recognized as forming beta-barrels at the time of prior searches and accounts for 20.8% of the increase in COB. We additionally identified previously uncharacterized BUL16 and BUL22 categories, which contribute 12.9% of the increase in COB.

**Figure 2.**
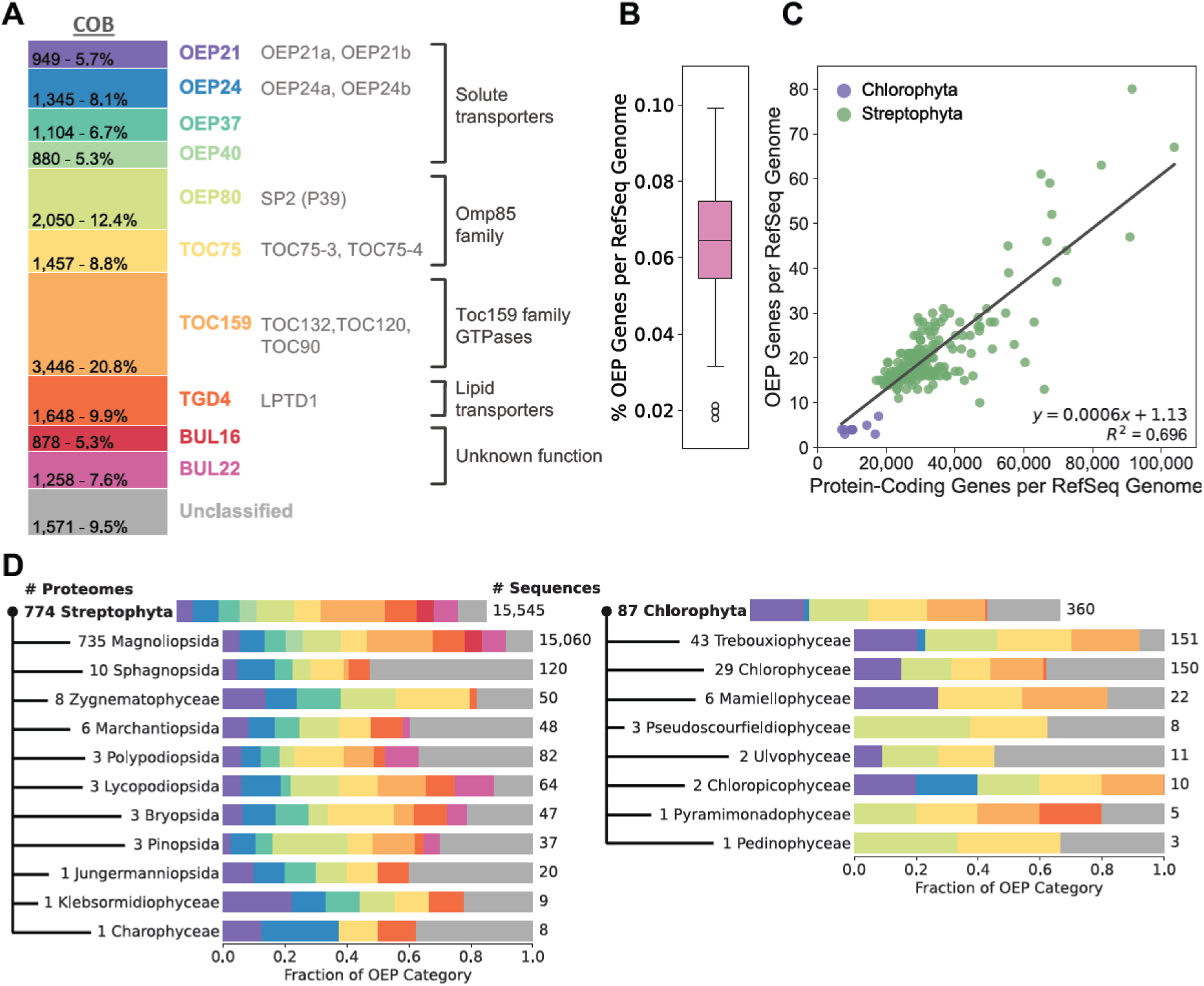
Features of COB. (A) Count and percentage of each OEP category within COB, along with homologous members included in each category, divided into broad functional groups. (B) OEP genes as a percentage of total protein-coding genes across 195 RefSeq genome assemblies. (C) Correlation between RefSeq genome size (protein-coding gene count) and OEP gene count per assembly for Streptophyta (green) and Chlorophyta (purple) species (R^2^ = 0.695). (D) OEP category composition across taxonomic classes within Streptophyta (left) and Chlorophyta (right), shown as the fraction of total OEP sequences belonging to each category. The number of proteomes (out of 861) represented per taxonomic class is indicated on the right.

COB contains 16,586 beta-barrel proteins, organized into ten OEP categories and an Unclassified group (Fig. 2A). Each category has different structural features, including strand numbers, IMS and cytoplasmic domain architecture, and open or closed barrel, and yet they all share the N-terminal strand to the left of the C-terminal strand when orienting the intermembrane space (IMS) downward. The ten categories in COB are fairly equivalent in representation within the database, ranging from approximately 4-21% of the dataset (Fig. 2A). These categories can be further divided into the functional groups: solute transporters (OEP21, OEP24, OEP37, OEP40), lipid transporters (TGD4, LPTD1), Omp85 family members (OEP80, TOC75, SP2), the TOC159 receptor family (TOC159, TOC120, TOC132, TOC90), and unknown function (BUL16, BUL22) (Fig. 2A). Chloroplast OEPs constitute a very small percentage of reference plant proteomes, approximately 0.06% (Fig. 2B), and the OEP gene count scales with the size of the plant genome (Fig. 2C). Notably, the phylum Chlorophyta has fewer OEP barrels than Streptophyta, even after accounting for the relative differences in the number of protein-coding genes. The distribution of OEP categories also differs among taxonomic classifications under Viridiplantae (Fig. 2D), demonstrating that not only does Chlorophyta have fewer OEP barrels than Streptophyta, but that it is especially deficient in the variety of solute transporters. While Streptophyta tends to have four separate types of solute transporters (OEP21, OEP24, OEP37, and OEP40), Chlorophyta often exclusively uses OEP21 with occasional instances of OEP24. Chlorophyta also lacks BUL22 or BUL16 (which is restricted to Magnoliopsida in Streptophyta) and has limited TGD4.

#### Chloroplast OEPs span across Viridiplantae

To characterize the abundance and distribution of chloroplast beta-barrel proteins within plant proteomes, we mapped COB sequences to their respective NCBI genome assemblies (the deposited genome and protein sequences for individual species or strains). This resulted in 861 assemblies, of which 195 were RefSeq genomes and used for further analysis. Mapping protein accessions to their NCBI gene identifiers and collapsing redundant entries allowed us to gather the number of unique OEP genes per RefSeq genome. These barrel gene counts were divided by the total number of protein-coding genes reported for each genome. These results show that chloroplast beta-barrel genes account for approximately 0.06% of the total plant genome (Fig. 2B). Additionally, the number of unique OEP genes detected per genome scales with overall proteome size (Fig. 2C, R^2^ = 0.695), suggesting the number of OEP genes expands proportionally with genome size across Viridiplantae. The number of OEPs show a wide range from 3-80 unique OEP genes in a given RefSeq genome. This likely reflects the expansion of gene families through duplication events in larger plant genomes. This is illustrated by *T. aestivum,* a hexaploidy genome with three ancestral subgenomes [51] and the largest RefSeq genome in our dataset with 103,793 protein-coding genes. *T. aestivum* contains 67 unique OEP genes spanning all categories, which is consistent with the scaling of OEP content with genome size.

To examine OEP category composition across Viridiplantae, we analyzed the protein sequences in COB across all 861 proteomes. The distribution of individual OEP categories varies across the taxonomic classifications (Fig. 2D). Within Streptophyta, all ten OEP categories along with Unclassified barrels show a consistent presence across the represented taxonomic classes, demonstrating broad conservation throughout land plants and their close relatives. In contrast, proteomes within Chlorophyta show a more refined OEP set, with taxonomic classes primarily restricted to OEP21, OEP80, TOC75, TOC159, and sequences within the Unclassified category, while OEP24, OEP37, OEP40, TGD4, BUL16, and BUL22 are largely absent or rare.

#### OEP strand numbers

The ten OEP categories display considerable diversity in beta-strand count. Representative AF3 predicted structures for each OEP category reflect this structural diversity, illustrating variation in barrel size and domain architecture (Fig. 3). To characterize this structural diversity across COB, we predicted the beta-strand count of seven of the OEP categories using PolarBearal [62], a beta-barrel strand count prediction algorithm developed by our group. However, three of the barrel categories (OEP80, TOC75, and TOC159) were found to have primarily open barrel topologies, which were intractable to PolarBearal annotation. Therefore, these categories were assigned strand counts via manual annotation. The remaining seven categories had enough closed barrels for PolarBearal strand count estimation. We identified a consensus closed strand count for each category, despite some variation among individual sequences (Table 1). Notably, our predicted strand counts differ from prior computational estimates but agree with all experimentally determined structures, giving confidence to our strand count predictions.

**Figure 3.**
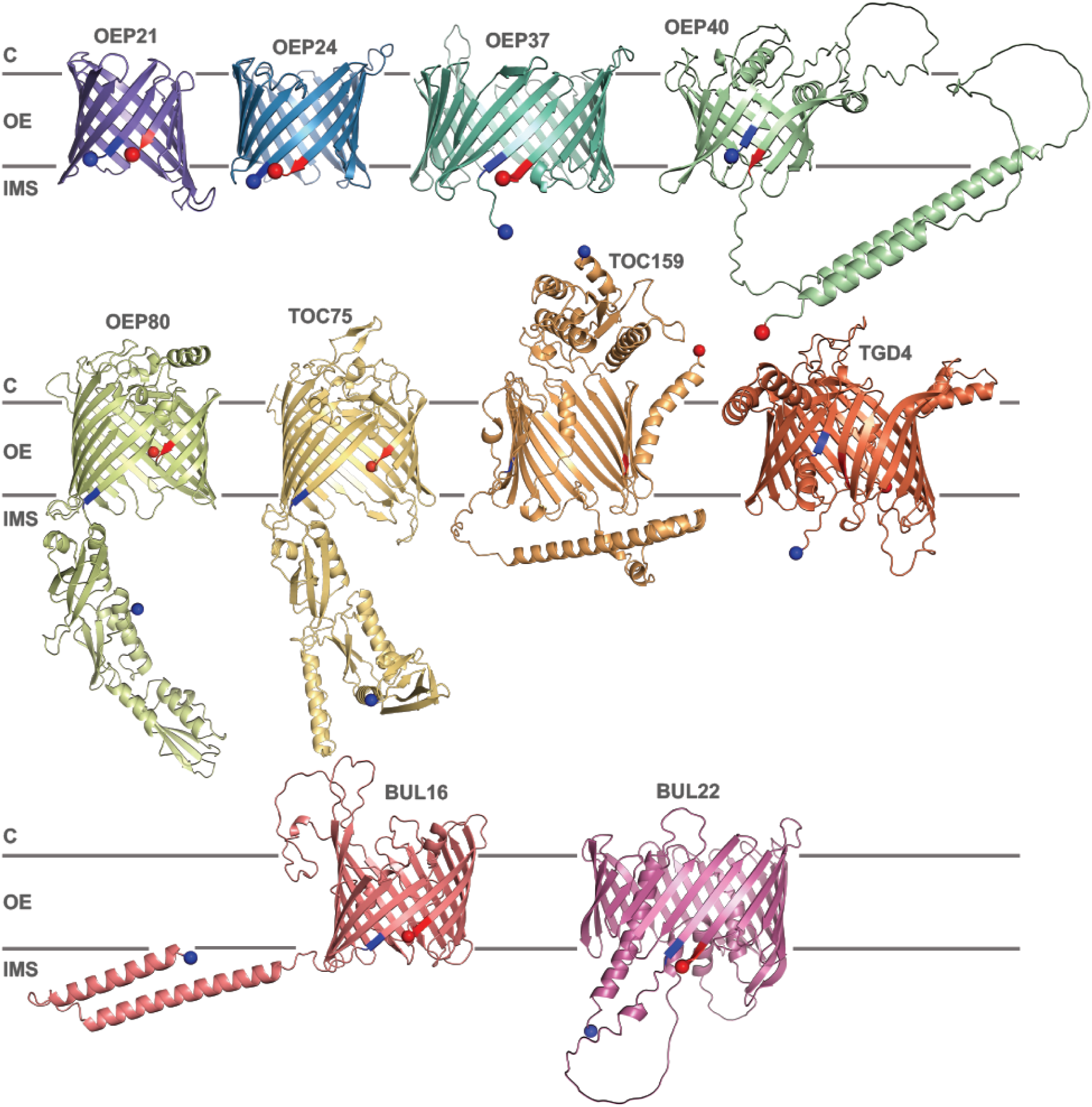
Representative AF3 structures for the OEP categories. OEP21 (GBG87084.1, purple), OEP24 (CAN8814779.1, blue), OEP37 (KAH6778313.1, teal), OEP40 (CAI0406690.1, green), OEP80 (CAG9465111.1, yellow-green), TOC75 (XP_020274341.1, yellow), TOC159 (CAE6140330.1, light orange), TGD4 (KAF3332909.1, dark orange), BUL16 (PIA58738.1, red), BUL22 (VAH09621.1, pink). Blue sphere = sequence N-terminus, red sphere = sequence C-terminus, blue residues = start residues for the first barrel beta-strand, red residues = end residues for the last barrel beta-strand. C = cytosol, OE = outer envelope, IMS = intermembrane space.

For porin OEP21, we detected a consensus of 12 beta-strands, consistent with the high-resolution NMR structure [10]. However, other categories demonstrated that previous literature consistently underestimated the strand numbers of OEPs. For the channel OEP24, prior estimates have ranged from 7 [13, 14] to 14 strands [43], while our analysis predicted a 16-stranded barrel. The glucose transporter OEP40 was previously predicted to be a 10-stranded beta-barrel [17, 43], while our analysis predicted 12 strands. Indeed, all successful PolarBearal predictions within the OEP40 category returned this count value, representing the highest consistency of any OEP category in our database. Unlike all other OEP categories with barrel domains located at the N-terminus, OEP40 positions its barrel domain at the C-terminus. It also has a variable length N-terminal helical region of predicted low structural quality. The OEP37 channel also shows an upward strand-count revision from previous predictions of either 12 [41], 16 [43], or 18 strands [63]. Our predictions indicate this category is dominantly 22-stranded barrels, the largest barrels in our dataset. Notably, these OEP37 barrels appear to lack a plug domain, a feature typically associated with bacterial porins of comparable size.

The lipid transporter TGD4 is structurally related to the bacterial lipopolysaccharide transport protein LptD, which forms a large 26-stranded beta-barrel [18, 20]. Previous studies have predicted TGD4 to consist of a 20-stranded beta-barrel [43], however, our analysis predicts TGD4 to be an 18-stranded barrel. This indicates that TGD4 represents a smaller version of the bacterial LptD fold. Interestingly, LptD is known to display an asymmetric barrel geometry, with a narrower diameter at one end and a wider diameter at the other (kidney bean shaped) [20]. We observe a similar pattern in TGD4, with one end of the barrel diameter appearing narrower than the other (Supp. Fig. 3).

When looking at our uncharacterized structures, two clusters appeared consistently throughout this set. The BUL22 category displays a 22-stranded barrel architecture that does not match any known OEP category (Fig. 3, Table 1). This structure contains an internal plug domain and adopts an asymmetric barrel with a wider diameter at one end and a narrower cobnstriction at the other (Supp Fig. 4). This is similar to the kidney bean shape of LptD previously discussed. The second prominent cluster, BUL16, displays a 16-stranded barrel topology and is also distinct from any characterized category. This structure has a variable length, low confidence N-terminal region (Fig. 3). These two clusters may represent novel eukaryotic beta-barrel classes that require further characterization.

**Figure 4.**
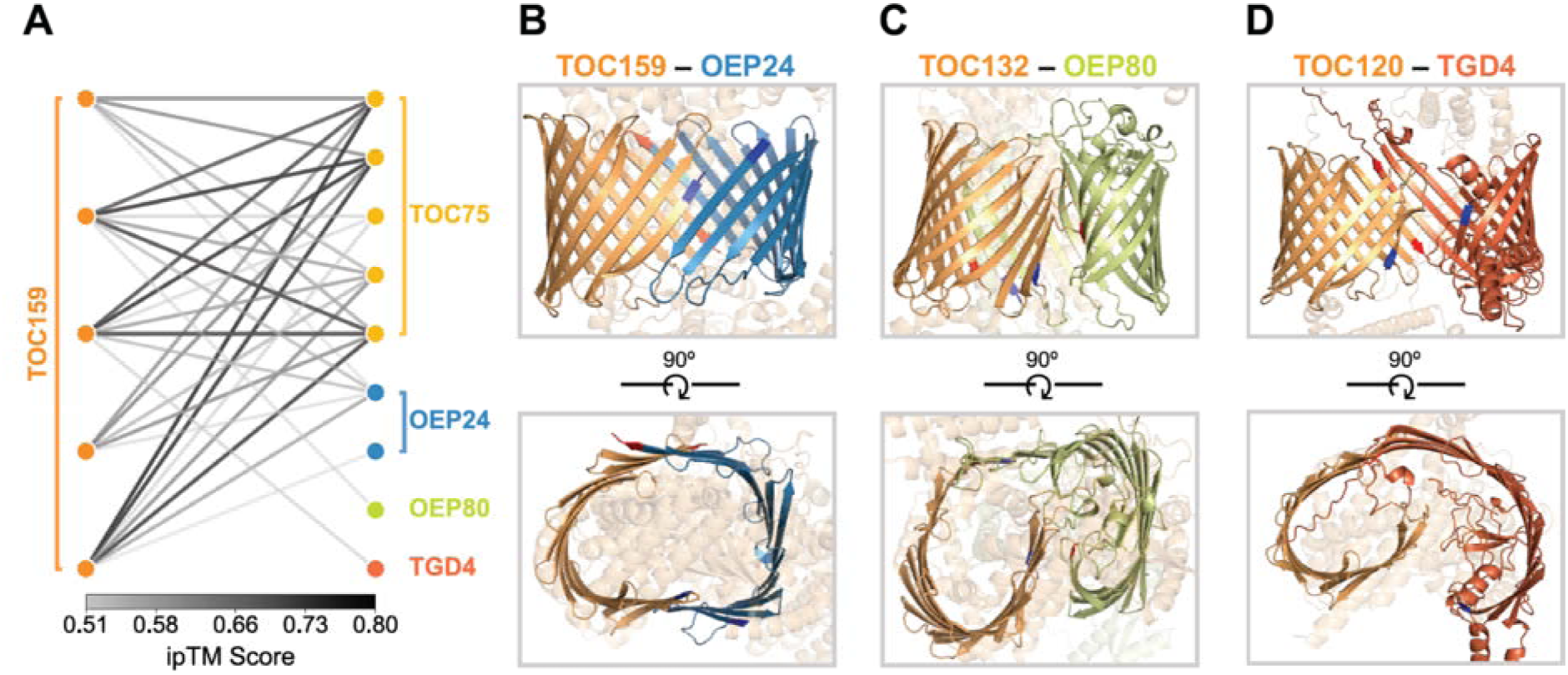
Predicted hybrid beta-barrel interactions in A. thaliana. (A) Graph network showing all manually verified hybrid barrel interactions from AF3 predictions. Each node represents a unique sequence in the specified OEP category. Edge darkness reflects ipTM score. (B) Top-scoring (ipTM = 0.63) hybrid barrel interaction between TOC159 (NCBI Protein Accessions: NP_567242.2, NP_001319848.1, NP_001328636.1, NP_001328635.1), and OEP24 (NP_001077680.1). (C) Top-scoring (ipTM = 0.52) hybrid barrel interaction between TOC132 (NCBI Protein Accessions: NP_179255.1, NP_001324729.1, NP_001324730.1), a member of the TOC159 family, and OEP80 (NP_568378.1). (D) Top-scoring (ipTM = 0.54) hybrid barrel interaction between TOC120 (NP_188284.1), a member of the TOC159 family, and TGD4 (NP_974244.1).

Three OEP categories, OEP80, TOC75, and TOC159, were almost exclusively open, yielding too few successful PolarBearal predictions for reliable strand count estimates (Supp. Table 2). Therefore, we randomly sampled 25 sequences from each of these three categories and manually assessed strand count using their AF3 structures (Table 1, Supp. Table 3). OEP80 and TOC75 were each found to consist of predominantly 16-stranded barrels, consistent with the conserved Omp85 superfamily architecture [64]. TOC159 was found to be a 14-stranded barrel. Both TOC75 and TOC159 strand number annotations are consistent with the cryo-EM structure of the TOC-TIC supercomplex [30, 31].

#### Open beta-barrel structures

Although to date, bacterial membrane barrels are exclusively known to be closed (where the first and last beta-strands interact), we observed that the structures of chloroplast barrels are both open and closed. However, strand counting with PolarBearal requires a closed barrel topology to generate a successful prediction. Therefore, we investigated whether PolarBearal success/failure was a reliable indicator of closed or open barrel topology. Using a manually curated test set, we found that PolarBearal failure predicts an open barrel with 94% accuracy (See Methods). We therefore used the rate of successful predictions within the non-redundant set of each OEP category as a proxy for the proportion of closed barrels within that category (Table 1). Seven categories, OEP21, OEP24, OEP37, OEP40, TGD4, BUL16, and BUL22 displayed a mixture of open and closed barrels, with open barrel percentages ranging from 26% to 68%. In contrast, OEP80, TOC75, and TOC159 were almost exclusively open, with OEP80 showing 96% open barrel conformations, and TOC75 and TOC159 were both 100% open (Table 1). Open barrel structures varied in their degree of openness, with some structures displaying only a marginal gap between the first and last beta-strands while others adopted a more prominent C-shaped conformation.

#### Lack of detectable beta-signal motif

A beta-signal motif has been proposed to reside in the last beta-strand of substrate proteins and is thought to be recognized by OEP80 during insertion into the outer envelope [24]. To investigate whether a conserved beta-signal is detectable across chloroplast OEPs, we searched for 6-10 residue sequence motifs using STREME and MEME [65]. These searches were performed on the non-redundant set of each OEP category, using both the full protein sequences and the last 50 residues separately.

While statistically significant hits were found for each category, none were consistently localized to the C-terminal beta-strand, and no motif spanned multiple OEP categories. This potentially reflects the high sequence similarity within OEP categories, even within our non-redundant sets at 70% sequence identity, rather than a definitive absence of a beta-signal. We also performed STREME analysis on all main OEP categories combined, excluding BUL and Unclassified sequences, at both 40% and 70% sequence similarity cutoffs. At 70% similarity, the analysis returned statistically significant motifs but with no consistency across individual motif searches, while at 40% similarity no significant motifs were detected. While there is biochemical evidence that demonstrates OEP80 interacting with the C-terminal beta-strands of substrate OEPs [24], it is also worth considering that our results could reflect biological diversity in substrate recognition. This could suggest that no universal C-terminal beta-signal exists across all OEP categories. Alternatively, multiple distinct motifs may be used across or within categories, or the beta-signal may be distributed elsewhere along the protein rather than in the last strand. A more sensitive analysis may require broader sequence diversity within each category or alternative approaches to motif detection.

### Candidate Hybrid Beta-Barrel Structures

One possible explanation for the preponderance of open barrel structures is that these structures are homomeric or heteromeric dimers that form a barrel together, i.e. hybrid barrel structures. TOC75 is known to form a hybrid beta-barrel with members of the TOC159 receptor family within the TOC complex, with structural studies demonstrating an interaction specifically between TOC75 and TOC90 [28, 30, 31]. This evidence, along with the observed high prevalence of open barrel topologies, supports the possibility that hybrid barrel formation may extend more broadly across chloroplast OEP categories. To investigate this possibility, we retrieved all 28 unique *A. thaliana* beta-barrel sequences from COB and performed pairwise AF3 predictions for each sequence against every other sequence, including self-interactions (homodimers). Predictions with an intermediate interface predicted template modeling (TM) score (ipTM ≥ 0.5) were retained, yielding 42 candidate interactions. Each was manually evaluated for hybrid barrel formation, and 31 were found to show hybrid barrel structures with contiguous interacting strands (Supp. Table 4). Notably, sequences within the TOC159 category were present in all 31 hybrid barrel interactions, indicating that TOC159 may serve as a universal interaction partner for hybrid barrel formation (Fig. 4A). These interactions involved not only TOC159 itself, but also other named proteins in the TOC159 category, supporting that hybrid barrel formation is potentially a property across the TOC159 family. The interacting partners included proteins in open barrel categories TOC75 and OEP80 as well as predominantly closed barrel categories OEP24 and TGD4 (Fig. 4B-D). Also, while lowering the ipTM threshold below 0.5 reduces confidence in predictions, we observed that OEP24 displayed similar universal hybrid-barrel behavior in the range of 0.4-0.5 ipTM, where it interacted with TOC159, OEP37, TOC75, and a sequence within the Unclassified category. While the TOC75-TOC159 interaction is consistent with the known TOC complex structure as previously stated, the interactions involving OEP24, OEP80, and TGD4 represent putative novel hybrid barrel complexes not previously described. If these interactions are real, TOC159 proteins may have multiple partners that are engaged either permanently by different proteins or that the same TOC159 swaps to have a different partner at different times.

### Comparison of chloroplast and bacteria sequences and contact maps

Throughout the development of COB, we observed features that differentiate chloroplast from bacteria beta-barrels at both the sequence and contact map level. To characterize these differences, we compared chloroplast OEPs in COB against bacterial OMPs from IsItABarrel. At the sequence level, chloroplast and bacterial beta-barrels show statistically significant differences in amino acid compositions (Fig. 5A). Chloroplast OEPs are enriched in charged residues, particularly positively charged amino acids, and show higher propensity for most non-polar residues. However, the beta-branched amino acids tend to have lower propensity in chloroplast compared to bacteria. Additionally, chloroplast sequences generally show lower propensity for aromatic, polar, and small amino acids relative to bacterial OMPs. These compositional differences potentially reflect the distinct membrane environments, functions, GC-content, and evolutionary histories between chloroplast and bacteria beta-barrels.

**Figure 5.**
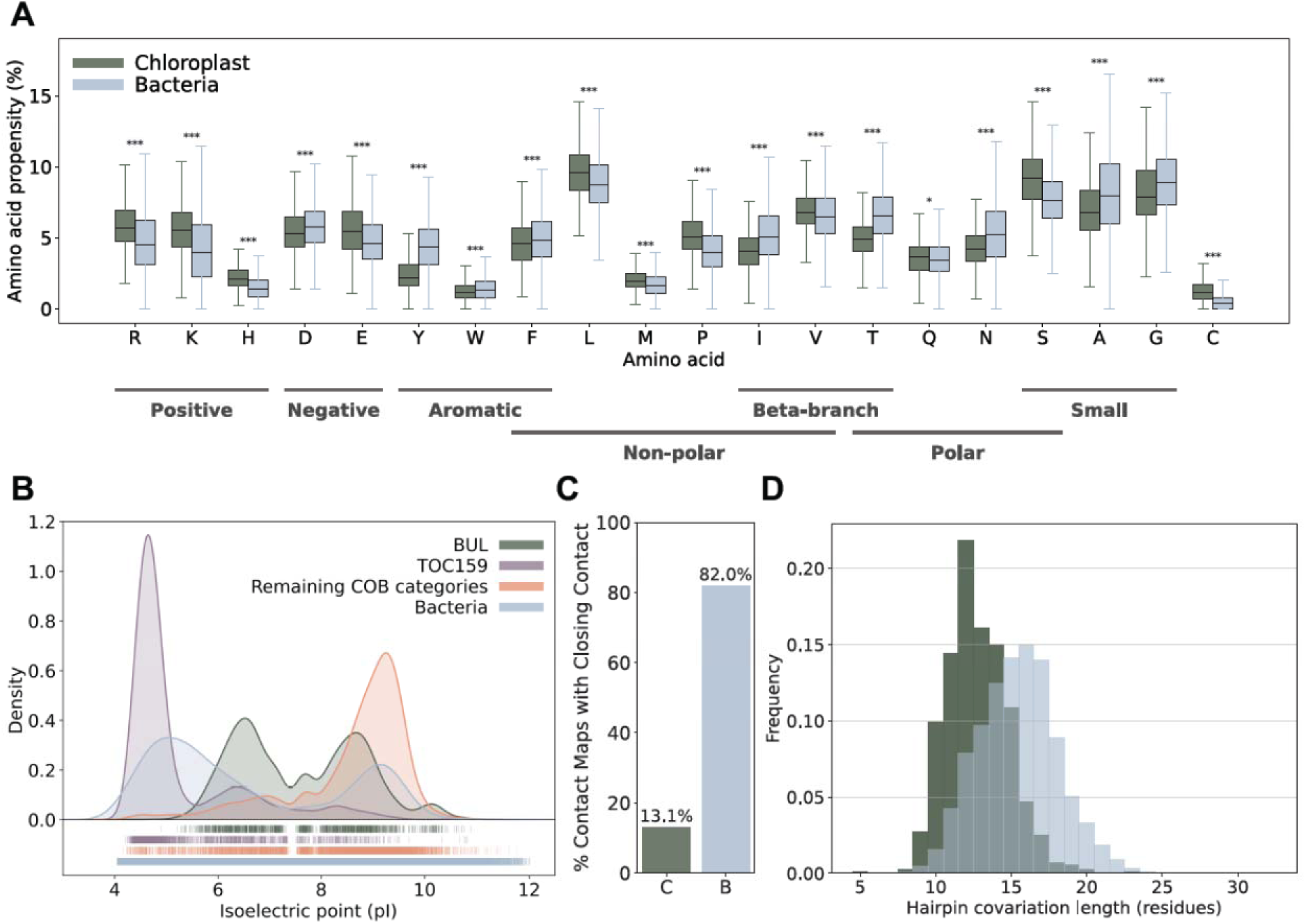
Comparison of chloroplast and bacterial outer membrane protein sequence and contact map properties. (A) Amino acid propensity for each of the 20 amino acids grouped by chemical property. Chloroplast OEPs are represented by a 40% non-redundant COB set (733 sequences) and bacterial OMPs by a previously constructed 30% non-redundant IsItABarrel set (101,152 sequences). In each boxplot, the box spans the 25th to 75th percentiles, and the center horizontal line indicates the median. Significance thresholds were defined as * p < 0.05, ** p < 0.01, and *** p < 0.001. Outliers were removed from the plot for visual clarity. (B) Kernel density estimate (KDE) of theoretical isoelectric point (pI) distributions for COB categories and bacterial OMPs, calculated from full protein sequences across the complete COB (16,586 sequences) and IsItABarrel (1,938,935 sequences) datasets. (C) Percentage of contact maps containing a detectable closing contact signal, determined by applying the bacterial IsItABarrel closing contact algorithm to the non-redundant COB set (733 sequences) and the non-redundant set of bacterial OMP contact maps generated in the IsItABarrel study (101,152 sequences). (D) Distribution of hairpin covariation lengths between the non-redundant COB set (733 sequences) and the non-redundant IsItABarrel set (101,152 sequences). Hairpin covariation lengths were collected using the COB classifier, and averaged for a given protein contact map.

Chloroplast OEP categories also differ from bacterial OMPs in theoretical isoelectric point (pI) values (Fig. 5B). As these values are calculated from full protein sequences, the pI of individual categories reflects contributions from all domains, not only the barrel itself. This is most evident in the TOC159 category, which displays a notably lower pI relative to other OEP categories. This is consistent with its highly acidic N-terminal domain pulling the overall sequence pI down. The BUL16 and BUL22 categories display a tight bimodal pI distribution. This pattern is distinct from the remaining COB categories, which tend toward higher, more basic pI values. Bacterial OMPs also display a bimodal pI distribution, though it is broader than that observed in the BUL groups.

Additionally, we find that bacterial OMPs showed a closing contact in 82.0% of contact maps, whereas only 13.1% of chloroplast OEP contact maps contained a closing contact (Fig. 5C). To determine this, we applied the bacterial IsItABarrel algorithm to COB contact maps to determine closing contact presence. This significant difference is consistent with the high prevalence of open barrel conformations observed across chloroplast beta-barrels, and further supports that open barrel topology is a distinguishing feature of chloroplast OEPs relative to bacterial barrels.

Bacterial OMPs also displayed longer hairpin covariation lengths on average (15.05 residues) compared to chloroplast OEPs (12.80 residues) (Fig. 5D). Hairpin covariation length reflects the approximate number of residues from contiguous strands interacting to form a hairpin contact on the contact map. This provides a rough estimate of strand length, suggesting that bacterial beta-strands tend to be longer than those of chloroplast OEPs.

### Unusual architectures: single-chain multi-barrel and hybrid barrel

Through our process of identifying chloroplast proteins, we found two structures that defied expectations of known chloroplast barrels. The first represents a single-chain multi-barrel fold, while the second exhibits an unusual hybrid barrel architecture (Fig. 6).

**Figure 6.**
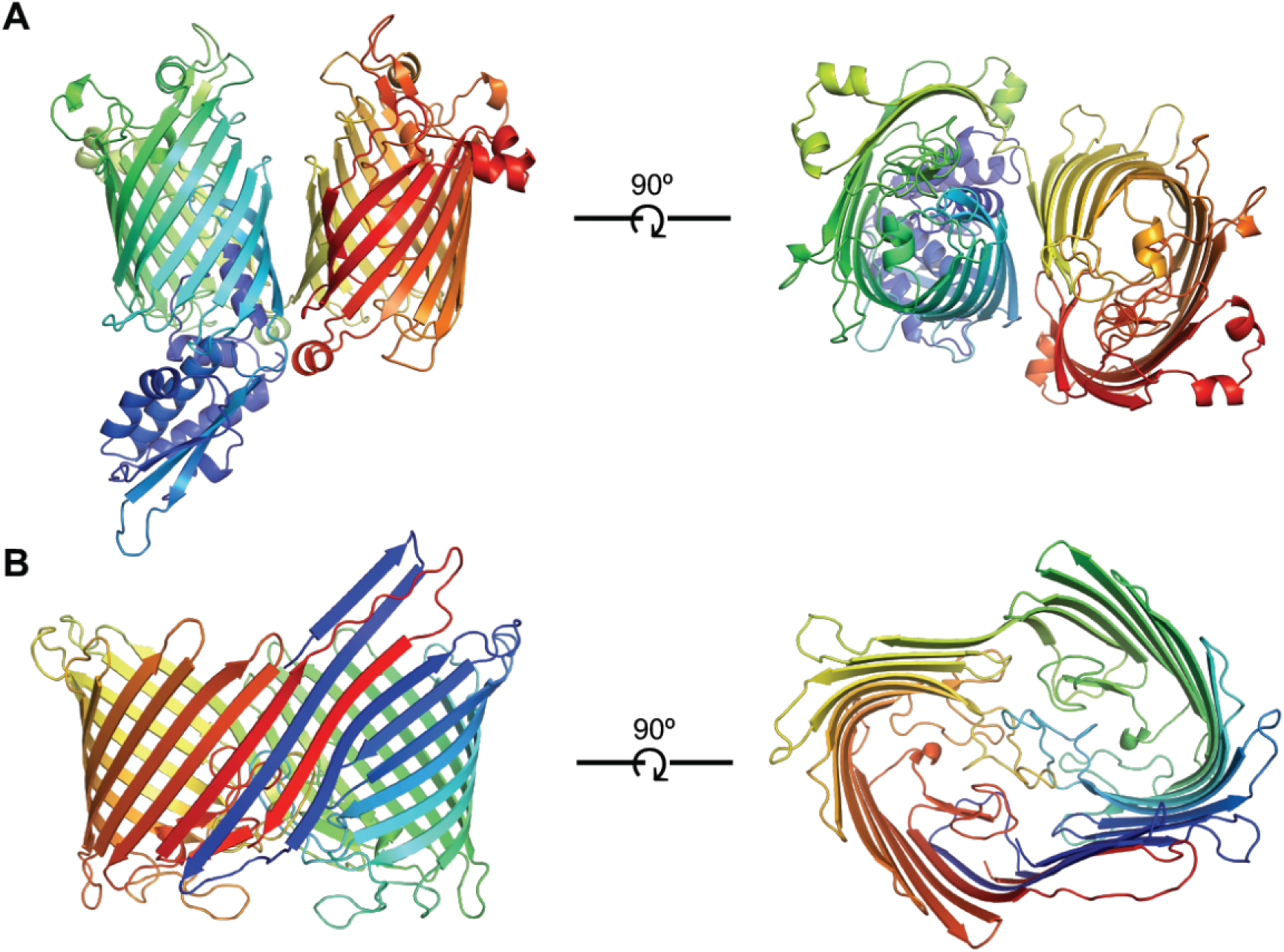
Single-chain beta-barrel architectures identified in COB. (A) Boltz-2 predicted structure of KAG7576803.1, a multi-barrel protein encoding two distinct barrel domains within a single polypeptide chain, with structural features consistent with OMP85-family proteins. (B) Boltz-2 predicted structure of KAM0881406.1, a single-chain barrel in which two barrel regions within the same polypeptide adopt a hybrid-like arrangement, classified within the OEP80 category.

Single-chain multi-barrel architectures have previously been described in Gram-negative bacteria [66]. During structural characterization of sequences within COB, we encountered two predictions displaying single-chain multi-barrel structures that have not been previously recognized in chloroplast outer envelope proteins. KAG7576803.1 resides in the Unclassified category, however, displays structural features consistent with OMP85-family proteins, including a singular POTRA domain and an identifiable gap between the first and last beta-strands (Fig. 6A). This architecture was reproduced in both AF3 and Boltz-2 [67] predictions.

An unusual hybrid-like barrel architecture was identified for KAM0881406.1 in the OEP80 category. Rather than adopting a continuous circular barrel, the Boltz-2 predicted structure displays a peanut-shaped structure in which two distinct barrel domains appear to share strands at their interface (Fig. 6B). This forms a larger, fused barrel from a single polypeptide chain. The AF3 prediction for this sequence also predicts two barrel domains within the same chain, with the open faces oriented towards each other, though without the continuous strand interweaving observed in the Boltz-2 structure (Supp. Fig. 5). Although the exact interactions between the two domains remain uncertain, both models agree on the presence of two-barrel domains within a single chain, supporting this as an interesting structure that calls for further structural and functional characterization.

## Discussion

In this study, we constructed COB, a comprehensive database dedicated to Chloroplast Outer envelope Beta-barrel proteins, consisting of 16,586 sequences across ten OEP categories and an uncharacterized group. COB was built using a machine learning classifier that identifies beta-barrel proteins based on features derived from evolutionary contact maps. This coverage allows characterization of chloroplast beta-barrels at a larger scale than was previously possible. It allowed us to perform structural analysis across entire OEP categories, quantify the prevalence of open beta-barrels, characterize the abundance and distribution of OEPs across plant proteomes, and identify candidate barrel interactions.

### Chloroplast beta-barrels are distinct from bacterial beta-barrels

Throughout the development of COB, we observed several differences between chloroplast and bacterial membrane beta-barrels. These differences are reflected in both the sequence and structural properties of the barrels, and informed the development of the COB classifier.

#### Chloroplast beta-barrels differ in amino acid preference compared to bacteria barrels

The distinct amino acid composition of chloroplast OEPs relative to bacterial OMPs may reflect differences in their respective membrane environments (Fig. 5A). In bacterial OMPs, aromatic residues form characteristic girdles at both membrane-water interfaces [9]. The reduced aromatic propensity in chloroplast OEPs may therefore reflect a reduced reliance on this anchoring mechanism. This is consistent with the fundamentally different lipid environment of the chloroplast OE. Unlike the bacterial OM, which is dominated by negatively charged lipopolysaccharide (LPS) in its outer leaflet, the chloroplast OE is enriched in galactolipids, which lack the charged headgroups. Additionally, we see an enrichment of positively charged residues in chloroplast OEPs (Fig. 5A), which may also be an effect of the lipid differences. In bacterial OMPs, a positive-outside rule has been described, where positively charged residues are enriched on the extracellular face of the barrel and interact with the negatively charged LPS in the outer leaflet [68]. The absence of LPS in the chloroplast OE suggests that the positive charge enrichment in chloroplast OEPs potentially serves a different purpose. Further, the differences in pI distributions across COB categories further supports that chloroplast OEPs are compositionally diverse in ways that may reflect their distinct domain architectures and function (Fig. 5B). The tight bimodal pI distribution observed in the BUL16 and BUL22 categories is notable, as it sets these categories apart from the remaining OEP categories. While the function of these groups remains unknown, this may indicate differences in their biology relative to the characterized OEP categories.

#### Chloroplast beta-barrels have shorter beta-strands than bacterial barrels

When evaluating the algorithm against chloroplast protein contact maps, we found that hairpin lengths were consistently shorter than those observed in bacterial OMP contact maps. To improve hairpin detection performance, we needed to lower the hairpin signal threshold and reduce the detection window size along the main diagonal of the contact map. To examine differences in hairpins more directly, we compared hairpin covariation lengths between the 40% clustered COB set and a non-redundant set of bacterial OMP contact maps from the IsItABarrel study (Fig. 5D) [48]. Chloroplast contact maps yielded an average hairpin covariation length of 12.80 residues, compared to 15.05 residues in bacterial contact maps. These values are measured in contact map pixels, where each pixel represents a single residue. This hairpin covariation length in the contact map represents interactions between the contiguous beta-strands, therefore, this can be interpreted as differences in the strand lengths across beta-barrels. This can generally be supported visually when comparing chloroplast OEP structural predictions with bacteria beta-barrels such as OmpA [69] or OmpT [70], that are generally taller and more elongated. While OmpA and OmpT are a small fraction of the types of beta-barrel structures found in bacteria, there is no comparable example to those beta-barrels characterized in chloroplast.

#### Limited sequence space of chloroplast OEPs compared to bacterial OMPs

Additionally, chloroplast contact maps showed more variability overall compared to those for bacterial OMPs. For instance, we observed two sequences that AF2 confidently predicted as membrane beta-barrels, yet their contact maps were essentially blank with no detectable features (Supp. Fig. 6). One potential explanation for this is the relatively lower number of sequenced homologs in chloroplast OEP families compared to bacterial OMP families. For instance, the bacterial phylum Proteobacteria (Pseudomonadota) (ncbitaxon:1224) has approximately 1.6 million genomes deposited in NCBI, whereas all of Viridiplantae (ncbitaxon: 33090) has only around 8,000 (Accessed April 2026) [71]. RaptorX contact prediction relies on coevolutionary information from multiple sequence alignments, and prediction accuracy scales with the number of available homologs [49]. Therefore, this could have been a potential explanation for the observed inconsistencies in reliable, detectable features. The reduced number of sequence homologs also restricted the size of the positive training dataset for the classifier, which ultimately was validated through manual evaluation of AF2 predicted structures alongside contact map features. This also played into the choice of the random forest model, because it is a relatively simple model and well suited to small datasets.

#### Chloroplast OEPs display open-barrel structures and form candidate hybrid barrels

While bacteria beta-barrels consistently display closed barrel topologies in which the first and last beta-strands interact, chloroplast beta-barrels appear to display a mixture of open and closed forms. Categories OEP80, TOC75, and TOC159 were found to be almost exclusively open, while the remaining categories, OEP21, OEP24, OEP37, OEP40, TGD4, BUL16, and BUL22 showed varying frequencies of each (Table 1). As these analyses are based on AF3 computational predictions, the high prevalence of open barrel conformations observed across OEP classes may reflect a systematic artifact of AF3 rather than true biological conformations. It is possible that some open barrel predictions may arise from incomplete sequences in the database, where a partial chain lacking one or more strands could prompt AF3 to predict an open conformation rather than a closed one. However, this pattern is consistent across all OEP classes and was not observed in analyses of bacterial beta-barrels. This mixture of open and closed conformations also manifested in the variation in the position and strength of the closing contact signal across chloroplast OEP contact maps compared to bacterial membrane barrels. This is supported by a direct comparison of closing contact prevalence between chloroplast OEPs and bacterial OMPs using the bacterial IsItABarrel closing contact algorithm [48]. Bacterial OMP contact maps contained a detectable closing contact signal in 82.0% of cases, compared to only 13.1% in chloroplast OEPs (Fig. 5C). However, because AF3 and RaptorX evolutionary contact maps both use MSAs, it’s unclear if these are both a feature of an MSA artifact, a feature of low information generating the MSA due to lack of proteomes, or a reality of protein structure in the chloroplast outer envelope.

If the preponderance of open barrel structures is not an MSA artifact, it is unclear if the high prevalence of open barrels across chloroplast OEPs is indicative of open barrels in the membrane or hybrid barrels in the membrane. It has been previously hypothesized that barrels need to be closed, as open barrels leave backbone carbonyls and amino groups exposed without satisfying hydrogen bonds. An open barrel position leaves one face exposed, which could allow it to pair with another open barrel to form a larger, hybrid assembly in which strands from two separate sequences complete a single continuous barrel. This is what occurs in the TOC complex, where TOC75 and TOC90 each contribute strands to form a hybrid barrel.

Here we found members of the TOC159 class (including the subclasses of TOC159: TOC120, TOC132, and TOC90) forming hybrid barrels with TOC75, OEP24, OEP80, and TGD4 (Fig. 4). It is notable that TOC159 category members adopted an open barrel conformation in 100% of the structures we evaluated, which may be directly related to its capacity for hybrid barrel formation (Table 1).

The TOC75-TOC159 interaction is well supported by experimental evidence and serves as an internal validation of our approach. The interactions involving TOC159 with OEP24, OEP80, and TGD4, however, represent putative novel hybrid barrel complexes with no prior characterization. If experimentally validated, these complexes could represent previously unrecognized assemblies with a range of novel functional roles.

Although OEP24 and TGD4 are predominantly closed barrel categories, they act as interaction partners with TOC159. This gives rise to the hypothesis that there may be some distribution of barrels where some are closed and some are in heterodimers. Another possibility is that some barrels transiently adopt an open conformation upon interaction and that such interaction is facilitated by something unique to the chloroplast outer envelope. Similarly, the formation of non-transient, heterodimeric, hybrid barrels in the chloroplast may require insertion mechanisms different from those for single domain barrels. A hybrid barrel consisting of two chains would likely require a more coordinated assembly pathway, potentially one that does not rely on C-terminal beta-signal recognition as proposed for canonical beta-barrels [24]. This may help explain why we do not see a C-terminal beta-signal in the proteins of COB.

### Strand count revisions

These analyses revealed upward revisions to strand counts across multiple OEP categories. For OEP24, prior estimates ranged from 7 to 14 strands, while our analysis predicts a 16-stranded barrel [13, 14, 43]. OEP40 was previously predicted to form a 10-stranded barrel [17, 43], whereas our analysis consistently predicts 12 strands. Also, OEP37 was predicted to form a 12 [41], 16 [43], or 18-stranded barrel [63], but we predict a 22-stranded barrel (Table 1). Earlier strand count predictions relied on sequence-based methods such as alternating hydrophobicity patterns and profile-based algorithms like PRED-TMBB2 [72]. These methods were developed based on bacterial OMPs, and not tuned for potential subtle differences between bacteria and chloroplast beta-barrels, such as shorter strand lengths. These prior strand count analyses have also largely been limited to individual sequences or small sets of proteins. By applying PolarBearal across larger sequence sets of each OEP category, we obtained consensus strand counts supported by hundreds of closed barrel predictions per family.

### OEP37 predictions lack an internal plug domain

OEP37 is predicted to form a 22-stranded beta-barrel, making it the largest characterized barrel in our dataset (Table 1). In bacterial OMPs, barrels of this size are consistently associated with plug domains. For instance, TonB-dependent transporters FhuA [73] and FepA [74] are well characterized examples, forming 22-stranded beta-barrels with an N-terminal plug domain that occupies the barrel interior. This architecture is conserved across all known TonB-dependent transporters [75]. Previous work has proposed that barrels larger than 18 strands may require a stabilizing plug domain to maintain structural integrity [76]. Therefore, the consistent absence of a plug domain in the predicted OEP37 structures is an unexpected finding. Additionally, the 26-stranded bacterial LptD barrel is plugged by a separate lipoprotein LptE, showing that this function can be carried out by a separate protein [20]. Whether something analogous exists for OEP37 is unknown, but the high-affinity blockage of OEP37 by the N-terminus of Tic32 suggests a protein interaction may contribute to its regulation or structure [16]. Whether OEP37 represents a genuine plug-free barrel or a separate protein fulfills this role, this is an area for future structural and functional studies.

### Chloroplast OEP composition varies across Viridiplantae

Chloroplast outer envelope beta-barrel genes only constitute approximately 0.06% of the total plant genome (Fig. 2B). It should also be noted that our percentages are calculated relative to the total plant genome, and normalizing to chloroplast-localized proteins only would likely give more comparable percentages to what has been seen in bacteria. To exemplify this comparison, we mapped the *A. thaliana* proteins in COB to their respective TAIR gene IDs [50], yielding 21 unique entries (Supp. Table 1). Given that ∼3,000 proteins are imported into the *A. thaliana* chloroplast [77], OEPs represent ∼0.7% of the chloroplast proteome. This is more comparable to the fraction in bacteria where beta-barrels account for approximately 0.27%-6.79% of the genome [48]. However, nearly all transmembrane proteins in the outer membrane of bacteria take on a beta-barrel fold, and the chloroplast outer membrane contains proteins with both alpha-helical and beta-barrel transmembrane domains, so the two systems have different protein fold compositions [8, 78].

Streptophyta and Chlorophyta represent the two major taxonomic lineages within Viridiplantae. Streptophyta encompass the charophyte algae and all land plants, while Chlorophyta comprise the morphologically diverse green algae [79, 80]. This biological distinction is potentially reflected in their OEP category compositions (Fig. 2D). Streptophyta retain all ten OEP categories consistently across their proteomes. In contrast, Chlorophyta contains a more refined set, with solute transporters OEP24, OEP37, OEP40, and lipid transporter TGD4 largely absent or rare. The retention of OEP21, OEP80, TOC75, and TOC159 across both phyla is consistent with conservation of core outer envelope functions required for metabolite exchange (OEP21) [11], protein import (TOC75/TOC159) [30], and outer membrane biogenesis (OEP80/SP2) across Viridiplantae [24, 25]. The absence of OEP24, OEP37, and OEP40 in Chlorophyta may reflect differences in metabolite trafficking between green algae and land plants. The relative abundance of OEPs already varies within land plants according to metabolite transport demand, as OEP24 and OEP37 are more abundant in C4 plants, while OEP21 predominates in C3 plants [15]. This pattern supports that OEPs are tuned to match specific metabolic demands for particular species. Chlorophyta exhibit distinct metabolic organization compared to land plants, including differences in carbon fixation, photorespiration, and central carbon metabolism [81, 82], which may alter the transport requirements across the outer envelope. In land plants, the demands of supplying energy across diverse tissues and cell types may have driven selection for these additional channels, whereas in Chlorophyta, OEP21 export may be sufficient. Additionally, the charophyte algae classes found in COB (Zygnematophyceae, Klebsormidiophyceae, and Charophyceae) also retain mostly all OEP categories (Fig. 2D). This pattern supports that diversification of the chloroplast outer envelope complement is an early evolutionary feature of Streptophyta, predating the emergence of land plants.

Furthermore, the absence of TGD4 across Chlorophyta may reflect a divergence in the machinery supporting ER-to-chloroplast lipid trafficking relative to plants. In land plants, lipid precursors synthesized at the ER must cross the chloroplast envelope membranes. Specifically, TGD4 receives ER-derived phosphatidic acid at the outer envelope, and TGD5 connects TGD4 to the inner envelope TGD1/2/3 complex to complete the transfer across the intermembrane space [18, 83]. However, both TGD4 and TGD5 are seemingly absent from *Chlamydomonas*, while TGD1/2/3 orthologues are retained [84–86]. Some ER-to-plastid lipid transfer does occur in *Chlamydomonas* through the conserved TGD1/2/3 complex [84], but no outer envelope component equivalent to TGD4 has been identified [85]. This supports that the inner envelope import functions are conserved, whereas the outer envelope machinery has diverged. The broad absence of TGD4 across Chlorophyta in COB indicates this is not specific to *Chlamydomonas* but a shared feature of the phylum. The taxonomic distribution of OEPs potentially reflects both conserved core and lineage specific transport related functions at the chloroplast outer envelope.

### Chloroplast OEPs display atypical multi-domain beta-barrel structures

Previously, proteins with multiple beta-barrel domains have been described in Gram-negative bacteria [66]. We detected structurally analogous architectures in the chloroplast outer envelope, which to our knowledge have not been previously reported (Fig. 6). The emergence of these structures in chloroplast may reflect similar evolutionary mechanisms to those proposed in bacteria, including tandem gene duplication and domain fusion [76, 87]. However, some of the multi-barrel architectures observed in chloroplast may differ from those described in bacteria. The high proportion of open barrel topologies in chloroplast OEPs, as described above, may permit intra-chain hybrid-like arrangements not seen in bacteria. This is exemplified by KAM0881406.1, where two barrel domains within the same chain appear to interact through their open faces (Fig. 6B). This type of architecture has not been described in bacteria, where barrels are predominantly closed. This indicates that open barrel topology may introduce additional possibilities for how multi-domain barrel architectures can be arranged and potentially diversify in function.

## Conclusions

We developed a contact map and machine learning-based classifier to identify chloroplast outer envelope beta-barrel proteins across Viridiplantae. The resulting database, COB, contains 16,586 sequences corresponding to unique protein accessions in the NCBI non-redundant (nr) database. These sequences are organized into ten OEP categories, including OEP21, OEP24, OEP37, OEP40, OEP80, TOC75, TOC159, TGD4, BUL16, BUL22, and an Unclassified group. The categories display considerable structural diversity in strand number, domain architecture, and barrel topology, including a prevalence of open barrel conformations not observed in bacterial outer membrane beta-barrels. Consistent with this, we identified candidate hybrid barrel structures in *A. thaliana*, in which two open barrels interact to form a larger, continuous beta-barrel assembly. We also identified single-chain multi-domain barrel architectures in the chloroplast outer envelope, a topology previously described only in Gram-negative bacteria. The distribution of OEP categories differs between Streptophyta and Chlorophyta, suggesting distinct biological roles at the chloroplast outer envelope, particularly in metabolite transport and lipid trafficking. Collectively, COB provides a comprehensive sequence resource to support future characterization of chloroplast beta-barrels, and to further investigate the evolution and function of the chloroplast outer envelope across the plant kingdom.

## Methods

### Training dataset 1,537DS

The 1,537 dataset (1,537DS) was created with the aim of constructing a high-confidence, low-redundancy dataset of predicted chloroplast beta-barrels, suitable for machine learning model training, development and evaluation. The dataset comprises 537 positive (beta-barrel) and 1,000 negative (non-barrel) sequences.

To collect positive sequences, we searched the UniProtKB database (release 2202_03) using profile hidden Markov models (pHMMs) for eight chloroplast beta-barrel families, OEP21, OEP24, OEP37, OEP40, OEP80, TGD4, TOC75, and OEP23, via the HMMER web server hmmsearch application, restricted to Eukaryota [88, 89]. This returned 6,099 sequences. After removing exact duplicates due to the homologous TOC75 and OEP80 families and filtering out sequences from non-plant species, 4,801 sequences remained. These sequences were evaluated for beta-barrel topology using the IsItABarrel algorithm and manual verification of AF2 structure, yielding 1,613 beta-barrel sequences [47, 48]. OEP23 sequences were included in the initial pHMM searches but were subsequently removed during AF2 structural validation, as predicted structures did not support a beta-barrel fold. To build a non-redundant positive set, we first clustered the 1,613 sequences at 75% identity using MMseqs2, producing a confident, core set of 349 representative sequences [54]. We then expanded this set by mapping the 1,613 sequences to UniRef50 clusters and retrieving 3,088 additional candidate sequences. These candidates were clustered at 75% identity, validated with AF2, and filtered to retain only sequences with less than 75% identity to any member of the core set of 349. This resulted in a final set of 537 positive sequences.

Negative sequences were drawn from NONTMBB1k, a previously curated set of 1,000 nonhomologous non-transmembrane beta-barrel proteins [48].

### COB classifier contact map feature extraction

Protein contact maps were generated using RaptorX [49]. Hairpins were detected by sliding a fixed sized window along the main diagonal of the contact map. Because beta-hairpins produce characteristic slashes perpendicular to the main diagonal, the sum of the anti-diagonal within each window position was used as a local signal for hairpin presence. Positions where this signal exceeded a threshold value were recorded as hairpin candidates. This is the same method described in the bacteria IsItABarrel algorithm for detecting hairpins, however with a lower threshold value of 9 [48].

The algorithm then locates the start and end coordinates of each hairpin by searching outward from the recorded hairpin position on the main diagonal. The search moves in two directions, both perpendicular to the main diagonal and a small range along the diagonal to account for positional variation. It evaluates a local window for the presence or absence of an anti-diagonal (hairpin) signal. The start coordinate is recorded where a filled anti-diagonal is first detected, indicating the onset of strand pairing, and the end coordinate where the signal transitions to empty, indicating the termination of the strand pairing.

Average hairpin length was defined as the mean Euclidean distance between the start and end coordinates across all detected hairpins. Hairpin spacing standard deviation was defined as the standard deviation of distances between consecutive hairpin positions along the main diagonal.

The closing contact score captures the characteristic anti-diagonal signal produced by interactions between the first and last strands of a closed barrel. To detect this, the algorithm identifies intersection regions between each set of hairpin pairs at least 3 hairpins away from each other. Within each candidate intersection, the anti-diagonal signal was evaluated by extracting a thick diagonal, formed by concatenating the main anti-diagonal and its two immediate neighbors. This thick diagonal was stepped incrementally in both directions along the main diagonal to account for slight variation in where the closing contact signal appears. The average value of this thick diagonal was computed at each offset position, and the maximum average across all offsets and all candidate intersections was returned as the closing contact score.

These features (number of hairpins, average hairpin length, hairpin spacing standard deviation, closing contact score, sequence length) were provided as input to the random forest classifier to produce a binary prediction of beta-barrel or non-beta-barrel.

### Random forest classifier training and evaluation

The 1,537DS dataset was split into training (90%) and test (10%) sets using a homology-based strategy to minimize sequence similarity between sets. Sequences were clustered at 30% identity using MMseqs2, and clusters were assigned entirely to one set, ensuring no closely related sequences appeared in both.

A random forest classifier was trained using scikit-learn with a grid search over five hyperparameters, including *bootstrap*, *class_weight, max_depth, min_samples_leaf,* and *min_samples_split* [90]. Model performance was assessed using 7-fold cross-validation. Folds were constructed from the same 30% similarity clusters, ensuring each fold contained sequences dissimilar to those in all other folds. Models were first filtered by retaining only those with cross-validation standard deviations of ≤ 0.12 across precision, Matthews Correlation Coefficient (MCC), and balanced accuracy. From the remaining candidates, the model with the highest combined MCC and precision was selected. Final performance was evaluated on the 10% held-out test set.

### COB creation

We classified all proteins from four selected genomes within the kingdom Viridiplantae. We downloaded the proteomes for *A. thaliana* (GCF_000001735.4), *C. reinhardtii* (GCF_000002595.2), and *T. aestivum* (GCF_018294505.1) from the NCBI RefSeq database (Accessed February 2024) [91]. The genome for the *Saccharum* hybrid cultivar R570 was downloaded from the Sugarcane Genome Hub (Accessed January 2025) [92]. Sequence annotations for Saccharum were later retrieved from Phytozome (Accessed September 2025) [93]. The proteomes were further processed, including implementing a length cutoff between 100 and 2,500 residues, removal of sequences with non-canonical amino acids, and removal of sequences with ‘mitochondria’ in the sequence header.

Each reference proteome was clustered at 40% sequence identity using MMseqs2. Taking the representatives from each cluster, we calculated the sequence similarity using EMBOSS Stretcher to any sequence already previously predicted by the COB classifier [94]. Representatives sharing ≥40% similarity with a previously evaluated sequence were assigned that sequence’s prediction directly, bypassing contact map generation. Representatives matching a previously evaluated negative sequence were discarded, whereas those matching at least one previously predicted positive sequence were retained as positives. In cases where a representative met the threshold to both positive and negative previously evaluated sequences, the positive assignment was retained. For those representative sequences that did not share ≥40% similarity with any previously evaluated sequence, contact maps were generated and evaluated by the COB classifier. Positive predictions were expanded to include all members of their respective clusters, and all resulting sequences were validated for beta-barrel topology using AF2.

Sequence annotations were retrieved from NCBI for *A. thaliana*, *T. aestivum*, and *C. reinhardtii*, and from Phytozome for *Saccharum* hybrid cultivar R570 (accessed September 2025) [93]. Sequences annotated as mitochondrial beta-barrels, including VDAC, Tom40, and Sam50, were removed. OEP40 and TOC159 were not recovered by the COB classifier workflow, therefore, these categories were supplemented using pHMM searches. The OEP40 pHMM used in the construction of the 1,537DS dataset and the Pfam model PF11886, corresponding to the TOC159 membrane domain, were searched against all four proteomes [43, 95]. Sequences recovered by the OEP40 and TOC159 pHMM searches were validated as beta-barrels by AF2. This resulted in a final high-confidence set of 387 chloroplast beta-barrel proteins, corresponding to 344 unique sequences.

#### Database expansion

The 344 unique sequences corresponding to 387 unique protein accessions were used as queries in a protein BLAST search against the NCBI non-redundant (nr) protein database, restricted to Viridiplantae (Accessed November 2025) [55, 56]. This search returned 30,286 unique candidate protein accessions. Pairwise sequence similarity between each of the 344 query sequences and the 30,286 candidates was calculated using EMBOSS Stretcher [94]. Only candidates sharing ≥40% sequence similarity with at least one query sequence were retained, resulting in 20,244 remaining protein accessions.

#### Annotation workflow

Each of the 20,244 candidate sequences was annotated either using its NCBI sequence description, or the annotation description provided by Phytozome for the *Saccharum* hybrid cultivar R570 sequences [93]. Descriptions were scanned using a set of keyword and pattern matching rules to identify candidate functional annotations across all known OEP categories, mitochondrial barrel proteins, and general barrel-related terms (porin, pore, barrel, channel, translocase, translocon, outer membrane, outer envelope). Each matched annotation was assigned a priority score, where specific category assignments such as OEP21 or TOC75 received higher priority than general terms such as “barrel” or “uncharacterized.” The highest priority annotation across all matches was selected as the final annotation for each sequence. There were three special cases that were handled. First, sequences matching two or more mitochondrial protein annotations were classified as general “mitochondrial”. Second, sequences matching two or more chloroplast protein annotations were classified as general “chloroplast”. Lastly, sequences matching both categories were annotated as a general beta-barrel, without an organellular assignment. Sequences with no keyword matches were assigned no annotation.

#### MMSeqs2 cluster classification

The 20,244 annotated sequences were clustered at 40% sequence identity using MMseqs2. Each cluster was assigned to one of eleven annotation groups, OEP21, OEP24, OEP37, OEP40, OEP80, TOC75, TOC159, TGD4, mitochondrial barrels, non-barrel proteins, or Unclassified, based on the consensus annotation of its members. The fraction of sequences belonging to each annotation group was calculated for every cluster, and the cluster was assigned to the group with the largest fraction. In cases where Unclassified represented the largest fraction, the cluster was reassigned to the next largest annotation group provided that group constituted at least 5% of the cluster members. Clusters assigned to the same group were then combined. Clusters classified as “mitochondrial barrels” or “non-barrel proteins” were removed from the dataset.

#### Foldseek structural classification

Clusters remaining in the Unclassified group following MMseqs2 annotation were subjected to structure-based classification using Foldseek [61]. To serve as Foldseek query structures, a representative AF3 structure was selected for each of the eight chloroplast OEP categories and three mitochondrial barrel proteins (VDAC, Sam50, and Tom40), chosen by visual inspection to confirm a confidently predicted, closed barrel structure. Mitochondrial representatives were included because sequence-based annotation alone may not have captured all mitochondrial proteins, and structural similarity provides an additional filter. For each MMseqs2 cluster defined as Unclassified, sequences were reduced to 70% sequence similarity using EMBOSS Stretcher data (See Redundancy reduction algorithm in Methods), and the remaining sequences were collected as classification targets. Each of the eleven representative query structures were searched against all the non-redundant Unclassified sequences using Foldseek easy-search with TMalign global alignment. For each target sequence, only query hits that met the minimum thresholds of 0.6 average TMscore and 0.7 query and target coverage were retained.

Classification of each non-redundant Unclassified MMseqs2 cluster proceeded in three steps. First, a consensus hit was required, meaning a cluster was eligible for classification only if all non-redundant members with a significant Foldseek hit agreed on a single query category. Second, the required proportion of cluster members with a matching hit varied depending on whether successful PolarBearal strand count predictions were available for non-redundant sequences in that cluster. For clusters with at least one strand count prediction, the strand count was required to match the expected consensus closed strand count for the assigned category (Table 1), and at least 10% of the non-redundant cluster members were required to have a hit to that query. For clusters lacking any strand count predictions, the threshold was raised to require at least 50% of members to have a hit to the consensus query. Third, a special case was applied for VDAC and Tom40. Because these two mitochondrial proteins share a 19-stranded topology and frequently produced overlapping Foldseek hits, clusters receiving exactly two unique query hits corresponding to both VDAC and Tom40 were classified as mitochondrial under the same strand count and hit rate thresholds as described. Clusters that did not meet these criteria remained uncharacterized and were retained in the Unclassified category.

#### Sequence length cutoff

Sequences less than 125 residues in length were removed from the dataset, as these are unlikely to encode a complete beta-barrel domain.

### Redundancy reduction algorithm

Non-redundant sequence sets were generated using a greedy algorithm applied to all-vs-all pairwise similarity scores from EMBOSS Stretcher [94]. Sequences were first sorted by length in descending order. Each sequence was then evaluated against all sequences already retained in the non-redundant set. If it shared similarity at or above the specified threshold of 70% with any retained sequence, it was discarded. Due to the longest sequences being processed first, this approach preferentially retains the longest representative when two sequences exceed the similarity threshold.

### PolarBearal validation as an open or closed beta-barrel indicator

Sequences were randomly sampled from the non-redundant sets across all OEP categories, with representation from each category, and each was manually evaluated as open or closed based on visual inspection of the AF3 predicted structure. Sampling continued until a balanced set of 45 open and 45 closed barrel sequences was obtained. We predicted the 90 AF3 structures using PolarBearal, and prediction success or failure was compared against the manual structural assignments to assess PolarBearal classification accuracy [62].

### Genome assembly mapping and proteome analysis

Genome assembly identifiers associated with each NCBI protein accession in COB were retrieved using the NCBI Identical Protein Groups tool, mapping 15,712 out of the 16,586 sequences to 860 unique genome assemblies [56]. *Saccharum* hybrid cultivar R570 sequences were excluded from this mapping step as their source assembly was already known and the proteome was not hosted on NCBI. Manually including the *Saccharum* sequences brought the total to 15,905 sequences mapping to 861 unique genome assemblies. To analyze OEP abundance per genome, assemblies were further restricted to those available in NCBI RefSeq under Viridiplantae, resulting in 195 RefSeq genomes. For each of these genomes, protein accessions were mapped to their corresponding NCBI gene identifiers and redundant entries were combined to produce unique OEP genes per genome. The number of protein-coding genes for each RefSeq genome was collected from NCBI as well. These values were used to calculate OEP gene percentage across plant proteomes and examine how OEP gene content scales with overall genome size (Fig. 2B-C). Analysis of OEP category distribution across taxonomic groups was performed using all 861 identified proteomes (Fig. 2D).

### Comparison of chloroplast OEPs and bacteria OMPs

Amino acid propensities were calculated from full-length sequences for both a 40% MMseqs2 non-redundant set of COB (733 sequences) and a 30% non-redundant representative set of bacterial OMPs from IsItABarrel (101,152 sequences) [48]. Statistical significance was assessed using the Mann-Whitney U test. Theoretical isoelectric points (pI) were calculated from full length sequences using BioPython [96] for the complete COB (16,586 sequences) and IsItABarrel (1,938,935 sequences) datasets. For analyzing closing contact presence, contact maps were generated using RaptorX for the 733 non-redundant COB sequences. The IsItABarrel algorithm was run on these 733 contact maps. The COB classifier was not used for this analysis as no validated cutoff exists for its closing contact score. Instead, the IsItABarrel algorithm was used, as a binary cutoff had already been established in the original study. For bacterial OMPs, closing contact presence data was taken directly from the IsItABarrel study for the 101,152 representative contact maps [48]. To gather hairpin covariation lengths, contact map features were extracted using the COB classifier for the 733 COB non-redundant contact maps and the non-redundant set of 101,152 bacterial OMP contact maps. To obtain the hairpin covariation lengths in residues, the hairpin length was first calculated as previously described using the Euclidean distance between hairpin start and end coordinates. Since these coordinates are oriented exactly diagonally on the contact map, dividing this value by √2 converts it to the number of pixels along the hairpin axis. This pixel count represents the hairpin covariation length, where each pixel corresponds to a single residue.

### Candidate hybrid beta-barrel structure prediction

All *A. thaliana* (GCF_000001735.4) beta-barrel sequences within COB were retrieved, yielding 36 unique protein accessions. Identical sequences were collapsed, producing 28 unique sequences for analysis. Pairwise AF3 predictions were performed for all 28 sequences against each other, including self-interactions, resulting in 406 predictions generated using default AF3 settings. Predicted complexes were filtered to retain those with an ipTM score of ≥ 0.5. This threshold was chosen to avoid excluding potentially relevant interactions, as the known TOC75-TOC159 interaction was observed across a range of scores extending below this value. Predictions meeting this threshold were manually evaluated for hybrid barrel formation, defined as at least one continuous strand interaction between the two barrel domains without a gap.

### Website creation

The structures in the COB database are searchable and available for download at https://cob.ku.edu. A user can search the database by chloroplast group protein type, sequence length, and/or portions of organism’s taxonomy classification. The page for individual proteins has additional information for the consensus strand count for the protein’s group type, average pLDDT, genome assembly id, and a visualization of the protein (Boltz-2 [67]) with the structure colored based on residue confidence values. Individual predicted structures can be downloaded from the search results page or the individual protein page. Alternatively, the entire database can be downloaded from the Search page as a bzip2 compressed tar file. This file will contain a Tab-separated value data file containing all of the records, a license file, and a directory containing all of the predicted structures in mmCIF format.

## Data, Materials, and Software Availability

All 16,586 COB protein structures predicted by Boltz-2 [67] are available for visualization, searching, and download at https://cob.ku.edu/. The COB classifier and the random forest model developed in this study are openly available at https://gitlab.ku.edu/sluskylab/cob/cob_classifier. Any additional data are available upon request.

## Acknowledgments

This work was supported by the NSF MRI 2117449, NSF MFB 2226804 to JSGS and Kansas INBRE, P20 GM1031418 to EP.

## Supplemental Figures

**Supplemental Figure 1.**
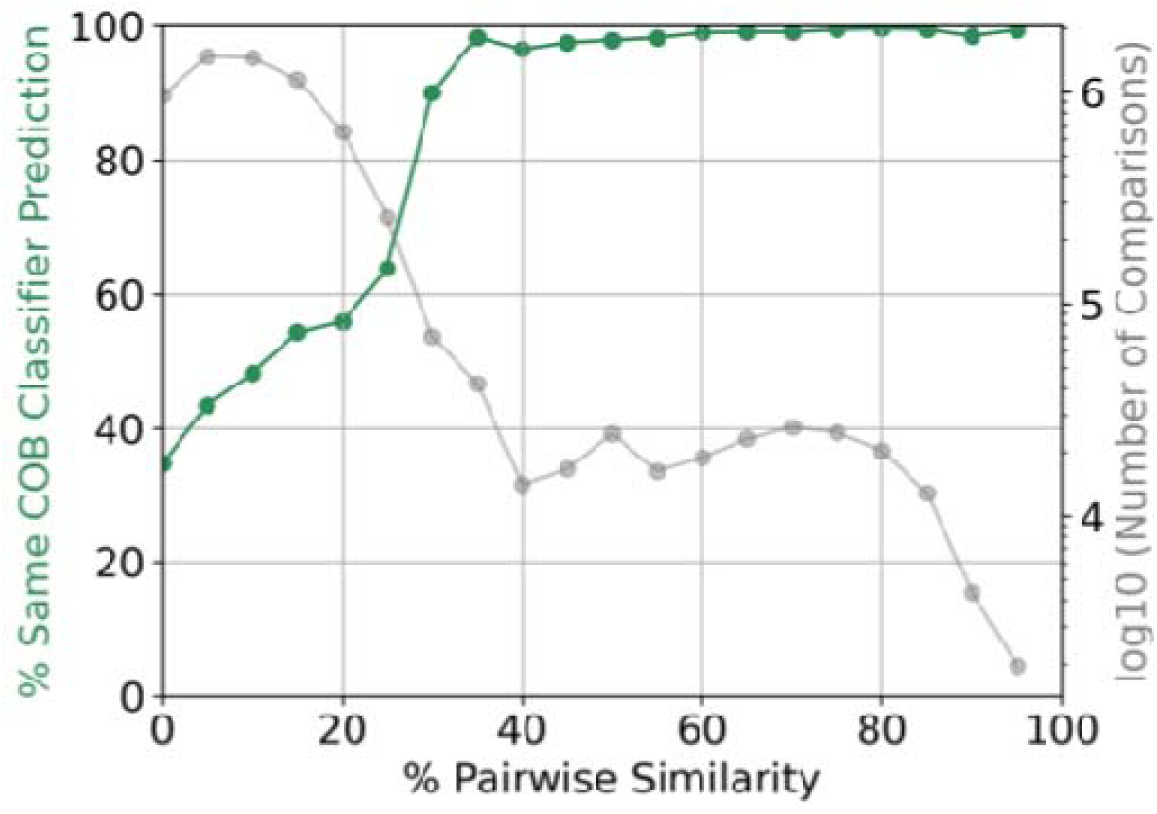
Relationship between pairwise sequence similarity and agreement in COB classifier predictions. Sequence pairs were binned by percent pairwise similarity, and the percentage of pairs within each bin receiving the same binary prediction is shown (green). The number of comparisons per bin is shown (gray). The dataset tested consisted of a balanced set of 1,770 sequences predicted positive and 1,770 predicted negative by the COB classifier. At ≥40% sequence similarity, greater than 97% of sequence pairs receive the same prediction, supporting the use of this threshold to bypass contact map generation for sequences similar to a previously predicted sequence.

**Supplemental Figure 2.**
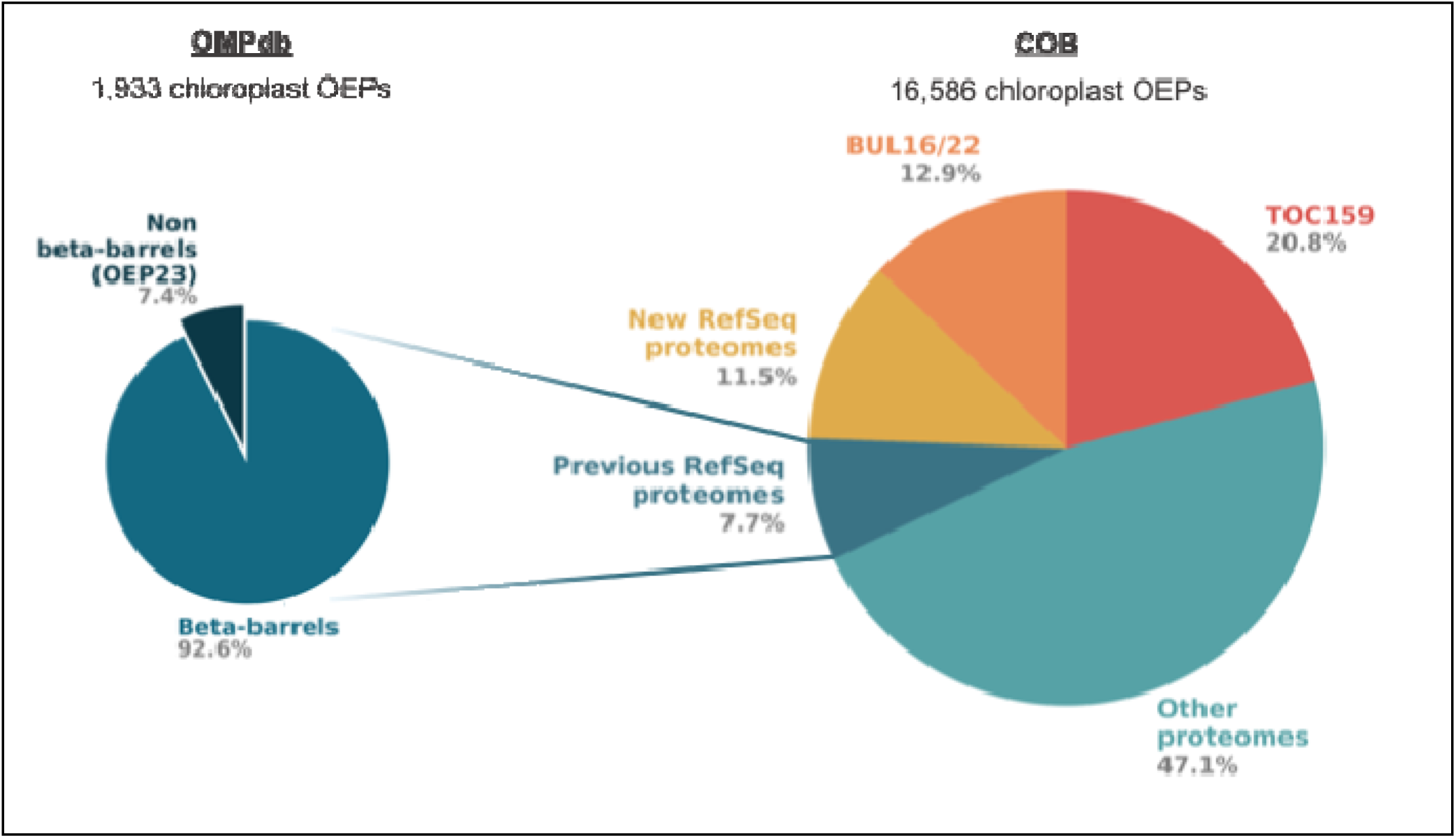
Comparison of OMPdb vs COB. OMPdb identified 1,933 chloroplast beta-barrels using the UniProt eukaryotic reference proteomes released in 2020, while COB contains 16,586 sequences. OEP23 structural predictions have since been shown to be inconsistent with a beta-barrel fold, and are placed in a non-beta-barrel section. COB sequences are partitioned by the source of increase from OMPdb chloroplast beta-barrels. Previous RefSeq proteomes are Viridiplantae NCBI RefSeq assemblies available in 2020 or before, approximating the proteome space accessible to OMPdb, and account for 7.7% of COB. New RefSeq proteomes are those released after 2020 and account for an additional 11.5%. Other proteomes are non-RefSeq assemblies excluded by OMPdb’s reference proteome restriction, accounting for 47.1%. The TOC159 receptor family was not represented in OMPdb and incorporation accounts for 20.8% of COB. BUL16 and BUL22 are previously uncharacterized beta-barrel categories identified in this work and contribute a further 12.9%.

**Supplemental Figure 3.**
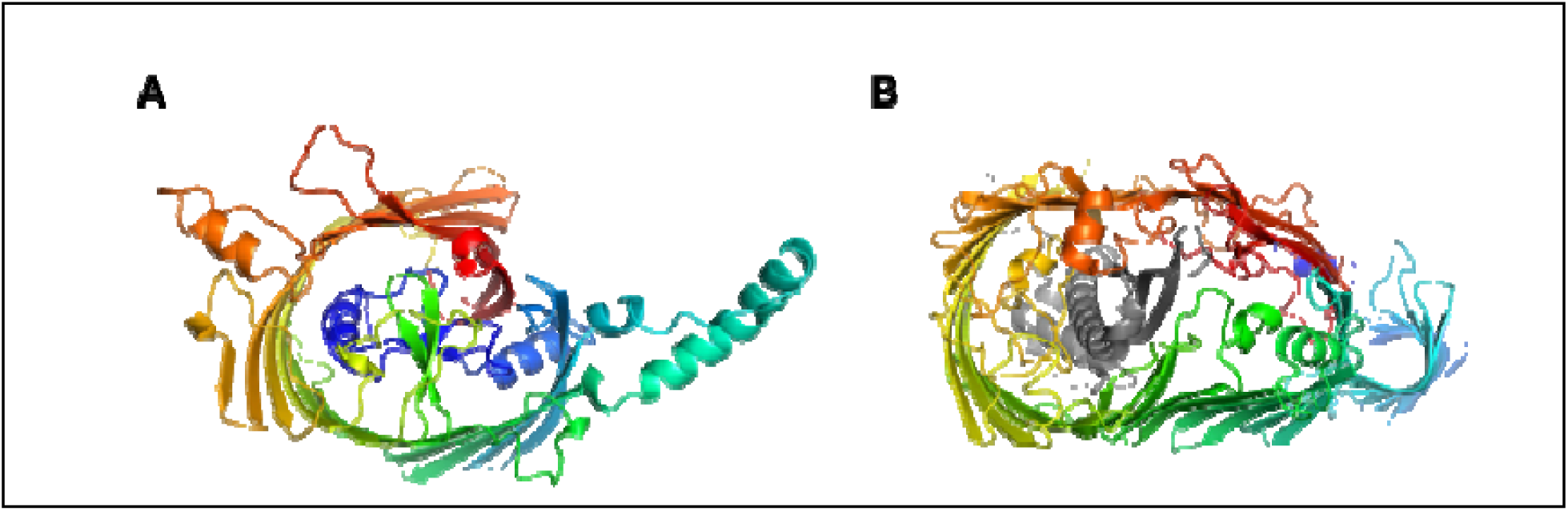
Structural comparison of TGD4 and the bacterial lipopolysaccharide transporter LptD. (A) Top-view of the AF3 predicted structure of XP_002982768.1, a representative TGD4 category sequence, showing an 18 stranded barrel with asymmetric geometry and a narrower diameter at one end. (B) Top-view of the crystal structure of LptD (PDB: 5IV9, rainbow) with its plug protein LptE (gray), forming a 26-stranded barrel with asymmetric geometry.

**Supplemental Figure 4.**
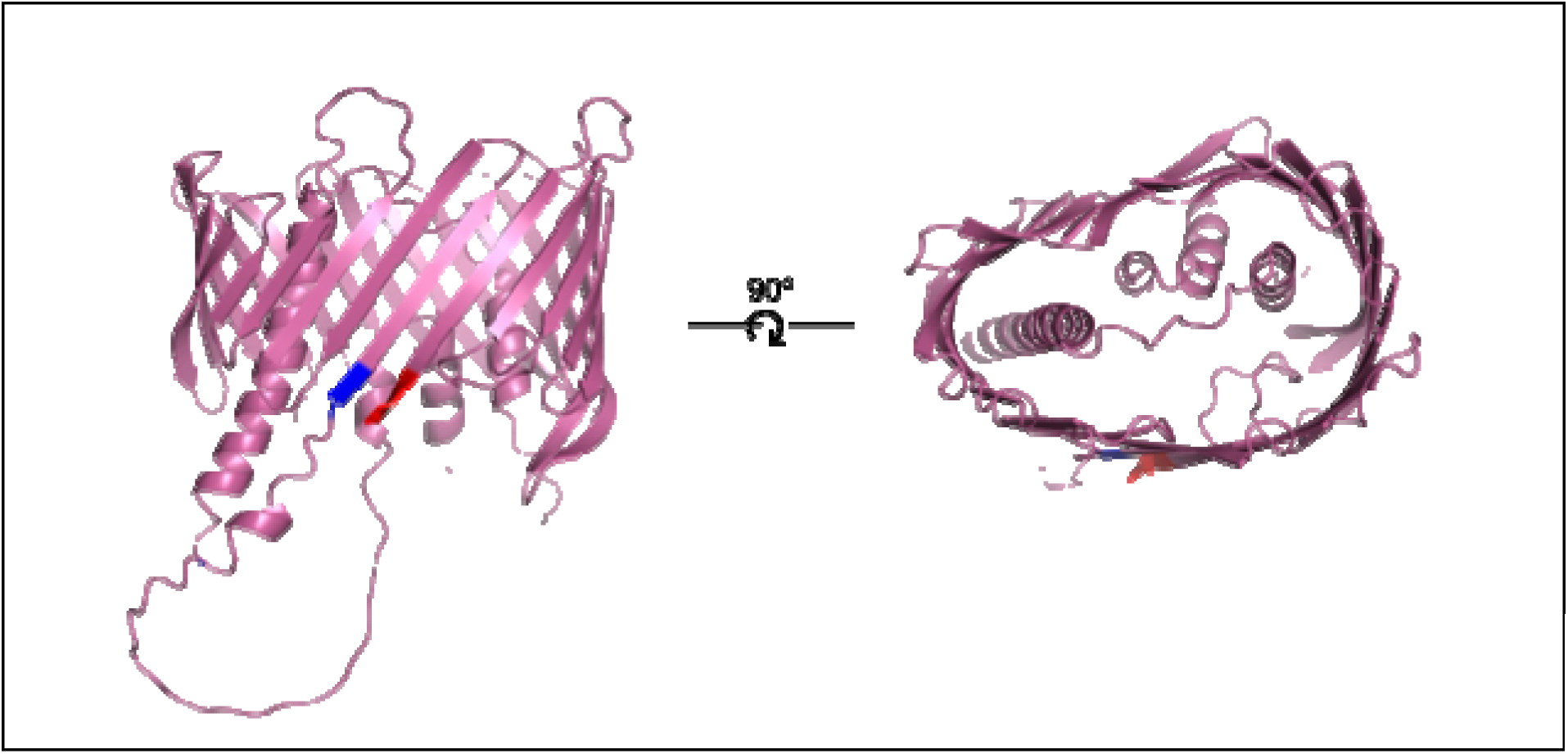
Side view (left) and top view (right) of the AF3 predicted structure of VAH09621.1, a representative BUL22 category sequence. The structure forms a 22-stranded barrel with an internal plug domain. The top view reveals an asymmetric, kidney bean shaped cross-section with a wider diameter at one end and a narrower diameter at the other.

**Supplemental Figure 5.**
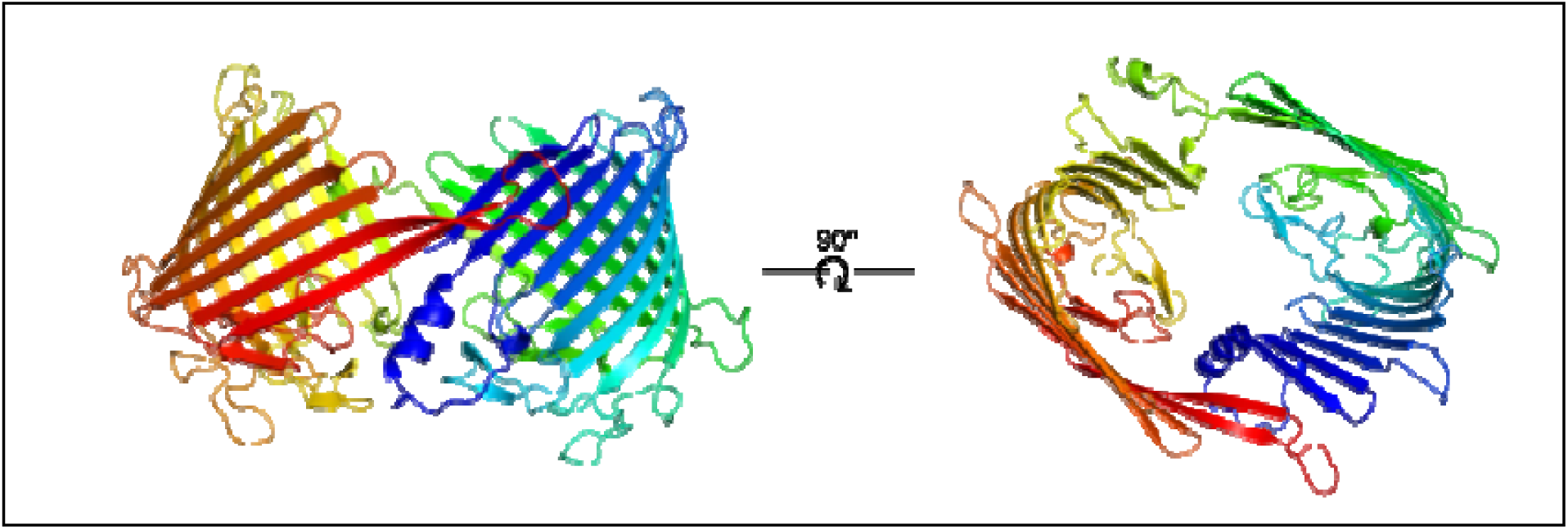
AF3 predicted structure of single-chain hybrid-like barrel, KAM0881406.1. The structure displays two barrel domains with the open faces facing each other.

**Supplemental Figure 6.**
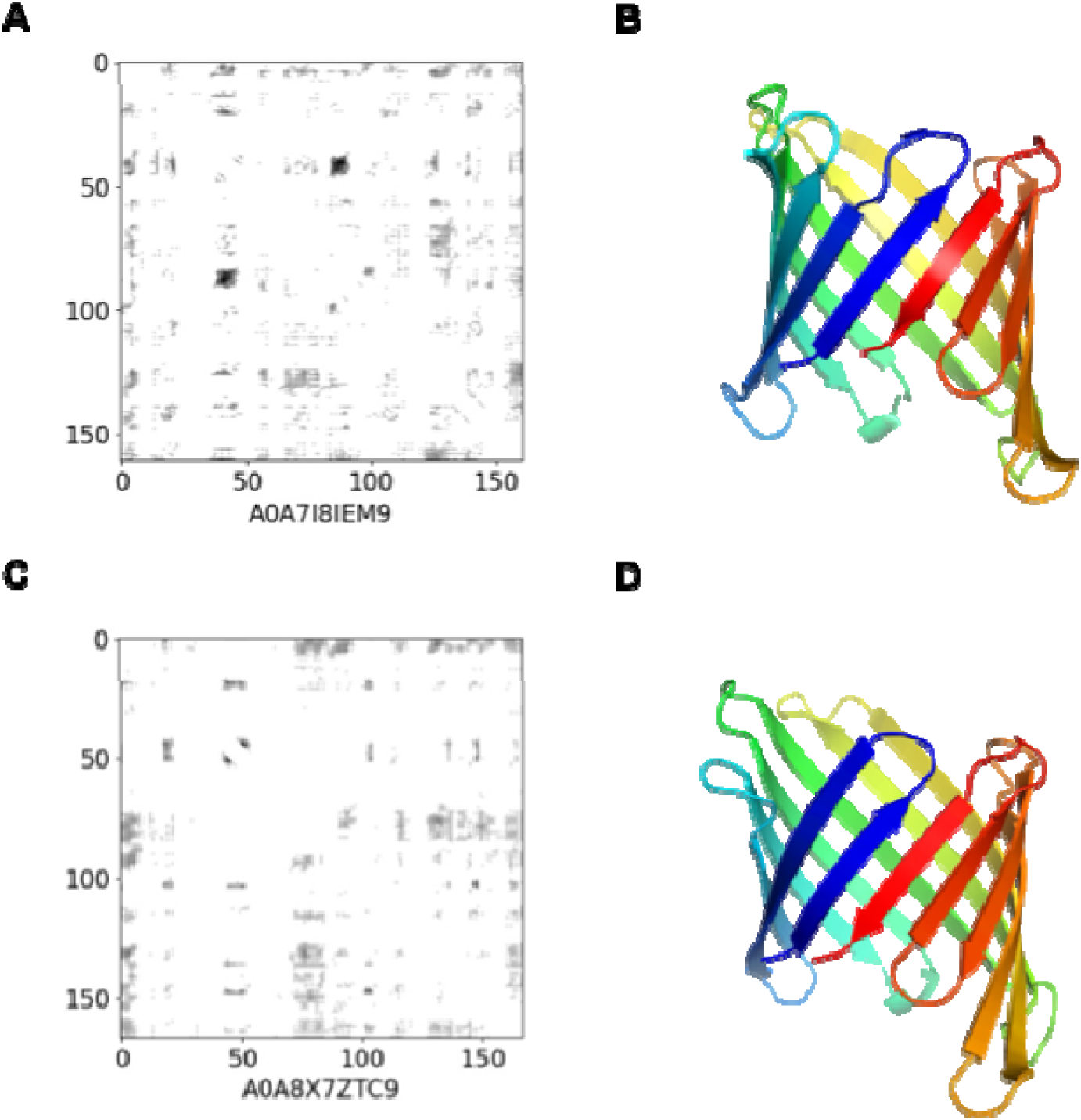
Examples of high-confidence AF2 structure predictions with low quality contact maps. (A) RaptorX contact map of A0A7I8IEM9 (UniProt). (B) AF2 structure prediction of A0A7I8IEM9 (pLDDT: 88.31), downloaded from the AlphaFold Protein Structure Database (AFDB) [97]. (C) RaptorX contact map of A0A8X7ZTC9 (UniProt). (D) AF2 structure prediction of A0A8X7ZTC9 (pLDDT: 91.25), downloaded from AlphaFold Protein Structure Database (AFDB).

**Supplemental Table 1.**
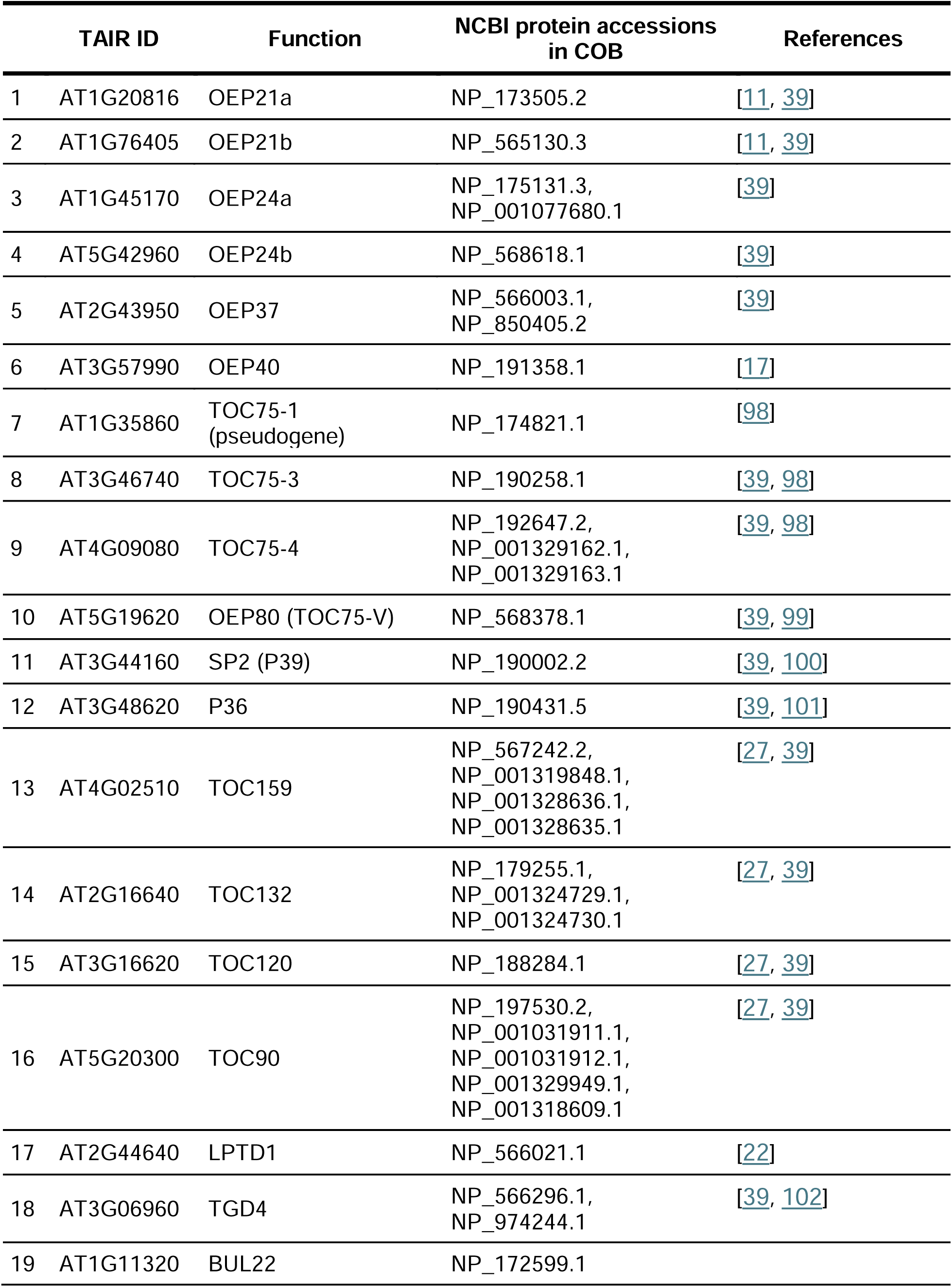
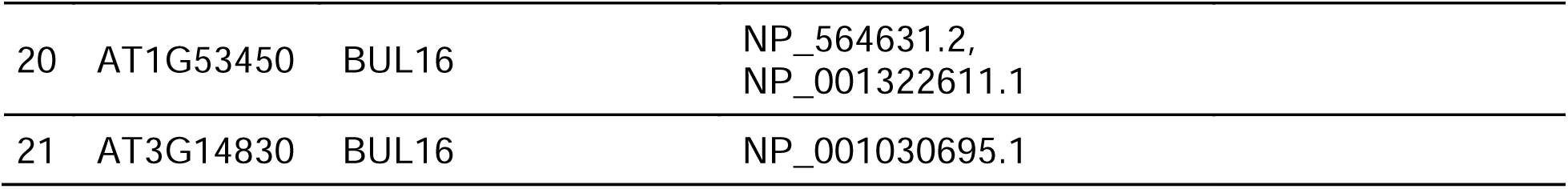
Mapping of Arabidopsis thaliana TAIR gene IDs to function and NCBI protein accessions in COB.

|  | TAIR ID | Function | NCBI protein accessions<br>in COB | References |
| --- | --- | --- | --- | --- |
| 1 | AT1G20816 | OEP21a | NP_173505.2 | [11, 39] |
| 2 | AT1G76405 | OEP21b | NP_565130.3 | [11, 39] |
| 3 | AT1G45170 | OEP24a | NP_175131.3,<br>NP_001077680.1 | [39] |
| 4 | AT5G42960 | OEP24b | NP_568618.1 | [39] |
| 5 | AT2G43950 | OEP37 | NP_566003.1,<br>NP_850405.2 | [39] |
| 6 | AT3G57990 | OEP40 | NP_191358.1 | [17] |
| 7 | AT1G35860 | TOC75-1<br>(pseudogene) | NP_174821.1 | [98] |
| 8 | AT3G46740 | TOC75-3 | NP_190258.1 | [39, 98] |
| 9 | AT4G09080 | TOC75-4 | NP_192647.2,<br>NP_001329162.1,<br>NP_001329163.1 | [39, 98] |
| 10 | AT5G19620 | OEP80 (TOC75-V) | NP_568378.1 | [39, 99] |
| 11 | AT3G44160 | SP2 (P39) | NP_190002.2 | [39, 100] |
| 12 | AT3G48620 | P36 | NP_190431.5 | [39, 101] |
| 13 | AT4G02510 | TOC159 | NP_567242.2,<br>NP_001319848.1,<br>NP_001328636.1,<br>NP_001328635.1 | [27, 39] |
| 14 | AT2G16640 | TOC132 | NP_179255.1,<br>NP_001324729.1,<br>NP_001324730.1 | [27, 39] |
| 15 | AT3G16620 | TOC120 | NP_188284.1 | [27, 39] |
| 16 | AT5G20300 | TOC90 | NP_197530.2,<br>NP_001031911.1,<br>NP_001031912.1,<br>NP_001329949.1,<br>NP_001318609.1 | [27, 39] |
| 17 | AT2G44640 | LPTD1 | NP_566021.1 | [22] |
| 18 | AT3G06960 | TGD4 | NP_566296.1,<br>NP_974244.1 | [39, 102] |
| 19 | AT1G11320 | BUL22 | NP_172599.1 |  |

|  |  |  |  |
| --- | --- | --- | --- |
| 20 | AT1G53450 | BUL16 | NP_564631.2,<br>NP_001322611.1 |
| 21 | AT3G14830 | BUL16 | NP_001030695.1 |

**Supplemental Table 2.** Complete PolarBearal strand count predictions and open/closed barrel topology percentages for all OEP category sets within COB.

| Category | Number of Sequences | Number of Nonredundant Sequences (70%) | Number of PolarBearal Success | % Open (PolarBearal Fail / Total) | Strand Number (Count) – % of Total Success |
| --- | --- | --- | --- | --- | --- |
| **OEP21** | 949 | 138 | 77 | 44% | 12 (73) – 95%<br>10 (2) – 3%<br>14 (1) – 1%<br>11 (1) – 1% |
| **OEP24** | 1,345 | 149 | 48 | 68% | 16 (46) – 96%<br>18 (2) – 4% |
| **OEP37** | 1,104 | 117 | 42 | 64% | 22 (35) – 83%<br>26 (4) – 10%<br>18 (1) – 2%<br>20 (1) – 2%<br>23 (1) – 2% |
| **OEP40** | 880 | 117 | 81 | 31% | 12 (81) – 100% |
| **OEP80** | 2,050 | 163 | 12 | 93% | 16 (8) – 67%<br>3 (4) – 33% |
| **TOC75** | 1,457 | 120 | 2 | 98% | 16 (1) – 50%<br>3 (1) – 50% |
| **TOC159** | 3,446 | 457 | 1 | 99.8% | 12 (1) – 100% |
| **TGD4** | 1,648 | 205 | 69 | 66% | 18 (65) – 94%<br>16 (2) – 3%<br>19 (2) – 3% |
| **BUL16** | 878 | 101 | 75 | 26% | 16 (73) – 97%<br>14 (2) – 3% |
| **BUL22** | 1,258 | 91 | 41 | 55% | 22 (35) – 85%<br>20 (4) – 10%<br>18 (1) – 2%<br>16 (1) – 2% |
| **Unclassified** | 1,571 | 598 | 92 | 85% | 12 (31) – 34%<br>16 (28) – 30%<br>18 (22) – 24%<br>22 (5) – 5%<br>19 (2) – 2%<br>3 (2) – 2%<br>20 (1) – 1%<br>14 (1) – 1% |
| **TOTAL** | 16,586 | 2,256 | 540 | 76% |  |

**Supplemental Table 3.** Complete manual strand count and open/closed barrel topology assignments for the three predominantly open OEP categories for which PolarBearal could not provide reliable consensus strand counts.

| Category | Open / Total (% Open) | Strand Number (Count) – % of Total |
| --- | --- | --- |
| OEP80 | 24 / 25 (96%) | 16 (10) – 40% |
|  |  | 17 (4) – 16% |
|  |  | 15 (4) – 16% |
|  |  | 14 (4) – 16% |
|  |  | 12 (1) – 4% |
|  |  | 11 (1) – 4% |
|  |  | 10 (1) – 4% |
| TOC75 | 25 / 25 (100%) | 16 (16) – 64% |
|  |  | 17 (3) – 12% |
|  |  | 15 (3) – 12% |
|  |  | 13 (2) – 8% |
|  |  | 8 (1) – 4% |
| TOC159 | 25 / 25 (100%) | 14 (23) – 92% |
|  |  | 12 (1) – 4% |
|  |  | 10 (1) – 4% |

**Supplemental Table 4.** The 31 AF3 pairwise predictions confirmed as hybrid beta-barrel interactions in A. thaliana, from a filtered set with ipTM ≥ 0.5. NCBI protein accessions are provided for each interaction partner. Where multiple accessions are listed for a single partner, these represent the same sequence.

| NCBI Protein Accession 1 | NCBI Protein Accession 2 | ipTM score |
| --- | --- | --- |
| NP_175131.3 | NP_567242.2, NP_001319848.1,<br>NP_001328635.1, NP_001328636.1 | 0.51 |
| NP_001077680.1 | NP_567242.2, NP_001319848.1,<br>NP_001328635.1, NP_001328636.1 | 0.63 |
| NP_001077680.1 | NP_001318609.1, NP_001031911.1,<br>NP_001329949.1, NP_197530.2 | 0.52 |
| NP_001077680.1 | NP_188284.1 | 0.59 |
| NP_001031912.1 | NP_001077680.1 | 0.51 |
| NP_001077680.1 | NP_001324730.1, NP_001324729.1,<br>NP_179255.1 | 0.59 |
| NP_001324730.1,<br>NP_001324729.1, NP_179255.1 | NP_568378.1 | 0.52 |
| NP_188284.1 | NP_974244.1 | 0.54 |
| NP_174821.1 | NP_567242.2, NP_001319848.1,<br>NP_001328635.1, NP_001328636.1 | 0.54 |
| NP_001329163.1 | NP_567242.2, NP_001319848.1,<br>NP_001328635.1, NP_001328636.1 | 0.79 |
| NP_190258.1 | NP_567242.2, NP_001319848.1,<br>NP_001328635.1, NP_001328636.1 | 0.67 |
| NP_001329162.1 | NP_567242.2, NP_001319848.1,<br>NP_001328635.1, NP_001328636.1 | 0.80 |
| NP_192647.2 | NP_567242.2, NP_001319848.1,<br>NP_001328635.1, NP_001328636.1 | 0.78 |
| NP_001318609.1,<br>NP_001031911.1,<br>NP_001329949.1, NP_197530.2 | NP_190258.1 | 0.60 |
| NP_001318609.1,<br>NP_001031911.1,<br>NP_001329949.1, NP_197530.2 | NP_001329163.1 | 0.68 |
| NP_001318609.1,<br>NP_001031911.1,<br>NP_001329949.1, NP_197530.2 | NP_001329162.1 | 0.66 |
| NP_001318609.1,<br>NP_001031911.1,<br>NP_001329949.1, NP_197530.2 | NP_192647.2 | 0.67 |
| NP_001031912.1 | NP_190258.1 | 0.60 |
| NP_001031912.1 | NP_001329162.1 | 0.66 |
| NP_001031912.1 | NP_192647.2 | 0.62 |
| NP_001031912.1 | NP_001329163.1 | 0.67 |
| NP_188284.1 | NP_567242.2, NP_001319848.1,<br>NP_001328635.1, NP_001328636.1 | 0.64 |
| NP_001329162.1 | NP_567242.2, NP_001319848.1,<br>NP_001328635.1, NP_001328636.1 | 0.76 |
| NP_174821.1 | NP_567242.2, NP_001319848.1,<br>NP_001328635.1, NP_001328636.1 | 0.53 |
| NP_001329163.1 | NP_190258.1 | 0.79 |
| NP_188284.1 | NP_001329163.1 | 0.76 |
| NP_001324730.1,<br>NP_001324729.1, NP_179255.1 | NP_001329162.1 | 0.64 |
| NP_001324730.1,<br>NP_001324729.1, NP_179255.1 | NP_192647.2 | 0.75 |
| NP_001324730.1,<br>NP_001324729.1, NP_179255.1 | NP_190258.1 | 0.79 |
| NP_001324730.1,<br>NP_001324729.1, NP_179255.1 | NP_001329162.1 | 0.77 |
| NP_001324730.1,<br>NP_001324729.1, NP_179255.1 | NP_192647.2 | 0.54 |

